# Parvalbumin^+^ and Npas1^+^ Pallidal Neurons Have Distinct Circuit Topology and Function

**DOI:** 10.1101/2020.02.14.950006

**Authors:** Arin Pamukcu, Qiaoling Cui, Harry S. Xenias, Brianna L. Berceau, Elizabeth C. Augustine, Isabel Fan, Saivasudha Chalasani, Adam W. Hantman, Talia N. Lerner, Simina M. Boca, C. Savio Chan

## Abstract

The external globus pallidus (GPe) is a critical node within the basal ganglia circuit. Phasic changes in the activity of GPe neurons during movement and their alterations in Parkinson’s disease (PD) argue that the GPe is important in motor control. PV^+^ neurons and Npas1^+^ neurons are the two principal neuron classes in the GPe. The distinct electrophysiological properties and axonal projection patterns argue that these two neuron classes serve different roles in regulating motor output. However, the causal relationship between GPe neuron classes and movement remains to be established. Here, by using optogenetic approaches in mice (both males and females), we showed that PV^+^ neurons and Npas1^+^ neurons promoted and suppressed locomotion, respectively. Moreover, PV^+^ neurons and Npas1^+^ neurons are under different synaptic influences from the subthalamic nucleus (STN). Additionally, we found a selective weakening of STN inputs to PV^+^ neurons in the chronic 6-hydroxydopamine lesion model of PD. This finding reinforces the idea that the reciprocally connected GPe-STN network plays a key role in disease symptomatology and thus provides the basis for future circuit-based therapies.

**Significance Statement:** The external pallidum is a key, yet an understudied component of the basal ganglia. Neural activity in the pallidum goes awry in neurological diseases, such as Parkinson’s disease. While this strongly argues that the pallidum plays a critical role in motor control, it has been difficult to establish the causal relationship between pallidal activity and motor (dys)function. This was in part due to the cellular complexity of the pallidum. Here, we showed that the two principal neuron types in the pallidum have opposing roles in motor control. In addition, we described the differences in their synaptic influence. Importantly, our research provides new insights into the cellular and circuit mechanisms that explain the hypokinetic features of Parkinson’s disease.

## Introduction

The basal ganglia are involved in motor control and adaptive behavior (Mink and Thach, 1991b; DeLong and Wichmann, 2007; Graybiel, 2008; Pennartz et al., 2009; Redgrave et al., 2010; Ito and Doya, 2011; Nambu and Tachibana, 2014; Jahanshahi et al., 2015; Dudman and Krakauer, 2016; Mink, 2018; Cox and Witten, 2019; Klaus et al., 2019). By providing a wide projection to all structures within the basal ganglia, the external globus pallidus (GPe) is theorized to critically influence information processing within this macrocircuit (Albin et al., 1989; DeLong, 1990; Albin et al., 1995; Parent and Hazrati, 1995; Mink, 1996; Bergman et al., 1998; Smith et al., 1998; DeLong and Wichmann, 2007; Kita, 2007; Hegeman et al., 2016). Because of the complexity in cellular composition and axonal projection patterns (Sato et al., 2000b; Mastro et al., 2014; Abdi et al., 2015; Dodson et al., 2015; Hernandez et al., 2015; Fujiyama et al., 2016; Saunders et al., 2016; Abecassis et al., 2020), it has been challenging to demonstrate the causal roles of the GPe in motor control.

We and others established that the GPe contains two principal neuron classes distinguished by their expression of the calcium-binding protein parvalbumin (PV) or the transcription factor Npas1. They account for roughly 50% and 30% of the GPe neuron population, respectively (Flandin et al., 2010; Nobrega-Pereira et al., 2010; Abdi et al., 2015; Dodson et al., 2015; Hernandez et al., 2015; Hegeman et al., 2016; Abecassis et al., 2020). PV^+^ neurons and Npas1^+^ neurons have different firing characteristics and projection targets. Specifically, PV^+^ neurons exhibit higher firing rates and preferentially target the subthalamic nucleus (STN) and substantia nigra pars reticulata (SNr), whereas Npas1^+^ neurons have lower firing rates and preferentially target the dorsal striatum (dStr) (Abdi et al., 2015; Hernandez et al., 2015; Saunders et al., 2016; Mastro et al., 2017; Abecassis et al., 2020). These results suggest that PV^+^ neurons and Npas1^+^ neurons likely play unique functional roles. Though this concept is fundamental to the organization and operating principles of the basal ganglia, no direct evidence has been presented.

To fill this critical knowledge gap, here we used contemporary circuit tools to examine the functional roles and synaptic inputs of GPe neuron subpopulations in mice. In accordance with their distinct electrophysiological and circuit characteristics, we found that PV^+^ neurons and Npas1^+^ neurons have distinct functional roles—PV^+^ neurons promote locomotion and Npas1^+^ neurons suppress it. By examining the excitatory inputs to the GPe, we showed that STN input to the GPe is unique in its cell type-specificity and preferentially targets PV^+^ neurons over Npas1^+^ neurons. Abnormal activity in the STN-GPe network has been postulated to underlie the hypokinetic symptoms of Parkinson’s disease (PD); however, the cell and circuit basis for its emergence remain poorly understood. By harnessing a chronic unilateral 6-hydroxydopamine (6-OHDA) lesion model of PD, we showed that the STN-PV^+^ input is selectively weakened in the chronic 6-OHDA lesioned model of PD. *In vivo* optogenetic stimulation of PV^+^ neurons promoted locomotion and alleviated hypokinetic symptoms in 6-OHDA lesioned mice. These results demonstrate the causal role of PV^+^ neurons in PD symptomatology, thus providing the basis for circuit manipulations in treatments of PD.

## Materials and Methods

### Mice

All procedures were done in accordance with protocols approved by Northwestern University Institutional Animal Care and Use Committees and were in compliance with the National Institutes of Health Guide to the Care and Use of Laboratory Animals. Experiments were conducted with the following mouse lines: A2a-Cre (Adora2a-Cre BAC, MMRRC 031168), C57BL/6J (Jax 000664), D1-Cre (Drd1a-Cre BAC, MMRRC 029178), D2-eGFP (Drd2-eGFP BAC, MMRRC 000230), Lox-STOP-Lox(LSL)-tdTom (Ai14, Jax 007914), Npas1-Cre-tdTom (Npas1-Cre-tdTomato BAC, Jax 027718), PV-Cre (PV-ires-Cre, Jax 017320), PV-tdTom (PV-tdTomato BAC, Jax 027395). The Npas1-Cre-tdTom BAC mouse was generated in-house (Hernandez et al., 2015). PV-Cre was crossed with LSL-tdTom to generate PV-L-tdTom (PV-Cre;LSL-tdTom) (Abecassis et al., 2020). Only heterozygous and hemizygous mice were used throughout the study to minimize the potential alteration of the phenotypes in mice carrying the transgene alleles (Chan et al., 2012). Mice were group-housed in a 12 h light-dark cycle. Food and water were provided *ad libitum*. All mouse lines were maintained by backcrossing with C57BL/6J stock. The genotypes of all transgenic mice were determined by tail biopsy followed by PCR to identify the presence of the relevant transgenes. Both male and female mice were used in this study.

### Stereotaxic injections

Mice aged postnatal day 30–35 were anesthetized in an isoflurane induction chamber at 3–4% isoflurane and immobilized on a stereotaxic frame (David Kopf Instruments). Anesthesia was maintained using 1–2% isoflurane. The scalp was opened using a scalpel and a small craniotomy (1 mm diameter) was made with a dental drill (Osada). Adeno-associated viruses (AAVs) were infused with calibrated 5 µl glass pipettes (VWR) pulled to have a tip diameter of 3 µm. The injection needle was left *in situ* for 5–10 min following the end of the injection to maximize tissue retention of AAV and decrease capillary spread upon pipette withdrawal. Experiments were performed 4–6 weeks after stereotaxic surgeries. The locations of the targeted injections were visually inspected under epifluorescence microscopy in *ex vivo* slices or histologically verified *post hoc*.

For *in vivo* channelrhodopsin2 (ChR2) stimulation of PV^+^ neurons or Npas1^+^ neurons, 90 nl of AAV_9_.EF1a.DIO.hChR2(H134R)-eYFP.WRE was injected into the GPe of PV-Cre or Npas1-Cre-tdTom mice. Alternatively, AAV_9_.EF1a.DIO.rev.eYFP.WPRE was injected into the GPe as a viral control. For *in vivo Guillardia theta* anion channelrhodopsin2 (GtACR2) inhibition of the GPe, 90 nl of AAV_1_.hSyn1.SIO.stGtACR2.FusionRed was injected into the GPe of PV-Cre or Npas1-Cre-tdTom mice. For *in vivo* GtACR2 inhibition of the subthalamic nucleus (STN), 45 nl of AAV_1_.CKIIa.stGtACR2.FusionRed was injected into the STN. For *in vivo* ChR2 stimulation of direct-pathway and indirect-pathway spiny projection neurons (SPNs), 720 nl of AAV_9_.EF1a.DIO.hChR2(H134R)-eYFP.WRE was injected into the dStr of D1-Cre and A2a-Cre mice, respectively. All surgeries for *in vivo* experiments were performed bilaterally.

To explore STN-specific ChR2 expression via retrograde Cre delivery, transduction patterns of AAV_retro_.Cre were first examined in LSL-tdTom mice; 90 nl of AAV_retro_.Cre was injected into the SNr. For Cre recombinase-inducible expression (CreOn) of ChR2 in the STN, 90 nl of AAV_retro_.Cre was injected into the SNr of the C57BL/6J mice and 45 nl of AAV_9_.EF1a.DIO.ChR2(H134R).eYFP into the STN. For *ex vivo* electrophysiological recordings of STN inputs, 45 nl of AAV_9_.Syn.ChR2(H134R).eYFP was injected into the STN. For recordings of pedunculopontine nucleus (PPN) and parafascicular nucleus of the thalamus (PF Thal) glutamatergic inputs, 90 nl of AAV_9_.Syn.ChR2(H134R).eYFP was injected. A full listing of the viral constructs and titer information is available in Tables 1 and 2.

**Table 1.**
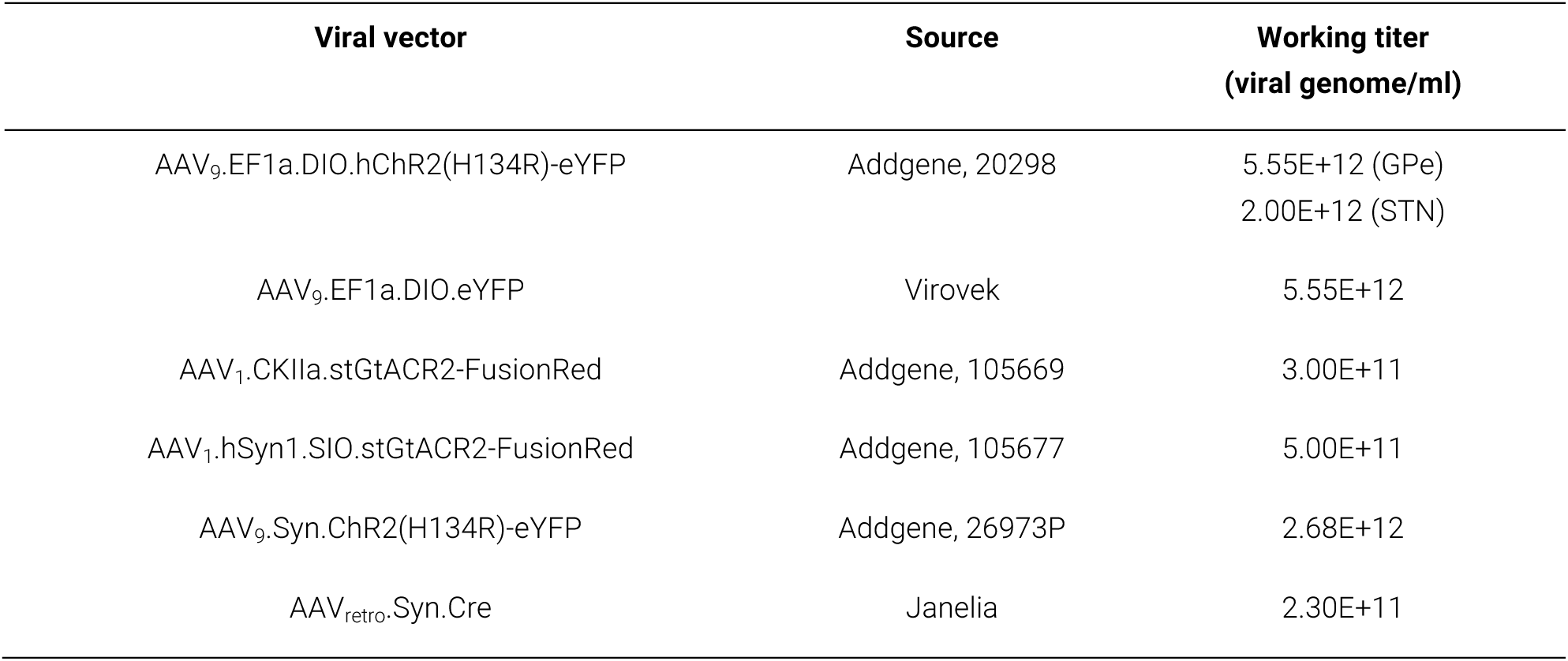
Detailed information on viral injections.

**Table 2.** Viral and 6-OHDA* injection coordinates.

| **Region** | **Volume per hemisphere (μl)** | **Rostral to Bregma (mm)** | **Lateral to Bregma (mm)** | **Ventral to the skull (mm)** |
| --- | --- | --- | --- | --- |
| GPe | 90 | −0.28 | 2.10 | 4.00 |
| STN | 45 | −1.70 | 1.60 | 4.50 |
| SNr | 90 | −3.00 | 1.60 | 4.20 |
| dStr | 720 | 0.90 | 1.40 | 3.20, 3.60 |
| PPN | 90 | −4.50 | 1.25 | 3.50 |
| PF Thal | 90 | −2.22 | 0.64 | 3.20 |
| MFB* | 990 | −0.70 | 1.10 | 4.95 |

Abbreviations: dStr = dorsal striatum, GPe = external globus pallidus, MFB = medial forebrain bundle, PF Thal = parafascicular nucleus of the thalamus, PPN = pedunculopontine nucleus, SNr = substantia nigra reticulata, STN = subthalamic nucleus.

### Fiber implantations and behavioral testing

Three weeks after stereotaxic injections, fiber optic cannulae were implanted bilaterally at the region of interest (Table 3). Each fiber cannula was assembled with an optical fiber (0.66 NA, 250 µm core) (Prizmatix) secured into a zirconia ferrule (1.25 mm O.D.) (Prizmatix) with non-fluorescent epoxy (Thorlabs). To determine the intensity of light exiting fibers, the output of fibers was measured with a power meter (Thorlabs). Cannulae were fixed to the skull using dental cement (Parkell). Mice were allowed to recover for 1–2 weeks before behavioral testing.

**Table 3.** Implantation coordinates.

| **Region** | **Rostral to Bregma (mm)** | **Lateral to Bregma (mm)** | **Ventral to the skull (mm)** |
| --- | --- | --- | --- |
| GPe | −0.28 | 2.1 | 3.8 |
| STN | −1.70 | 1.6 | 4.3 |
| SNr | −3.00 | 1.6 | 4.1 |
| dStr | 0.90 | 1.4 | 3.0 |

Abbreviations: dStr = dorsal striatum, GPe = external globus pallidus, SNr = substantia nigra reticulata, STN = subthalamic nucleus.

Behavioral tests were performed in a standard lit room between 2:00 pm and 7:00 pm. *In vivo* optogenetic experiments were performed in an opaque white plastic open-field box (28 cm × 28 cm), which was cleaned with 70% ethanol. On the day prior to optogenetic experiments, each mouse was allowed to acclimate to the open-field box for twenty minutes, with a fiber optic cable attached to the implant via a rotary joint (Prizmatix). Pre-trial, basal ambulatory activity was collected for five minutes at the beginning of the behavioral session. For ChR2 activation, mice were subjected to either a single, sustained light pulse or a patterned pulse train (5 ms pulses at 20 Hz) delivered for 10 s. For GtACR2 activation, only the sustained protocol was used. A blue (peak, ∼455 nm) LED (Prizmatix) was used to activate both opsins. The light power used for opsin activation was 12–18 mW measured at the fiber tip. Up to ten trials were run for any given protocol with a one-minute intertrial interval. Movement data were collected with an overhead camera at the maximum sampling rate (10 Hz). Mouse position was tracked using EthoVision XT (Noldus). After the completion of behavioral experiments, transgene expression patterns and fiber optic implant locations were histologically verified *post hoc*. Only histologically validated subjects were included in this study.

Speed of locomotion was computed from the distance traveled within the open-field arena per unit time. ‘Light-period’ corresponds to 10 s of light delivery. ‘Pre-period’ and ‘post-period’ correspond to the 10-s epoch before and after light delivery, respectively. Fold change in locomotor activity was calculated by dividing the difference in speed between light-period and pre-period by that of pre-period. Immobility was defined as periods with horizontal displacement less than or equal to 1 cm/s for at least 0.1 s. Mobility bouts were defined as periods with horizontal displacement more than 1 cm/s for at least 0.1 s. To calculate normalized speed, data for each trial were divided by the mean spontaneous locomotor activity measured from 25 s immediately before the light-period. The median differences in movement speed between pre-period and light-period and their 95% confidence intervals were estimated (https://www.estimationstats.com); five thousand bootstrap samples were taken; bias-corrected and accelerated corrections were applied to the resampling bootstrap distributions to adjust for both bias and the skewness (Ho et al., 2019). Logistic regressions were performed using R codes implemented in JASP (https://jasp-stats.org/). For single-trial analyses, data from both patterned and sustained stimuli were pooled. We considered logistic regressions for outcomes representing changes in speed during pre-period and light-period above the 0.1-, 1-, and 10-fold cutoffs for PV-Cre mice; or below the 0.1-, 0.5-, and 0.8-fold cutoffs for Npas1-Cre-tdTom mice, with each regression having the strain and manipulation condition as the explanatory variable. Speed during pre-period was considered as a covariate. To examine if there was hysteresis associated with optogenetic stimulation, trial number was considered as a factor in the model.

### Chronic 6-hydroxydopamine lesion

To study the changes in the STN-GPe network in a parkinsonian state, a chronic, unilateral 6-hydroxydopamine (6-OHDA) lesion model of PD was used to achieve a loss of dopamine neurons of the nigrostriatal system (Ungerstedt, 1968; Beal, 2001; Dauer and Przedborski, 2003; Meredith and Kang, 2006; Bezard and Przedborski, 2011). Unilateral 6-OHDA lesion was performed using standard procedures (Chan et al., 2011; Chan et al., 2012; Hernandez et al., 2015; Cui et al., 2016; Glajch et al., 2016). In brief, 6-OHDA (2.5 μg/μl dissolved in 0.9% w/v NaCl with 0.1% w/v ascorbic acid) was injected into the medial forebrain bundle (Table 2) of the left hemisphere. Three weeks after the 6-OHDA injection, lesion success was determined by performing the cylinder test to assess forelimb use impairments (Schallert et al., 2000; Iancu et al., 2005; Chan et al., 2011; Cui et al., 2016). The ratio of ipsilateral to total forepaw touches was used to determine the success of the lesion. In this task, during a five-minute exploratory behavioral assessment in a clear glass cylinder, weight-bearing contacts made by each forepaw on the glass walls of the cylinder were manually quantified. Forelimb use asymmetry was determined by calculating left, right, and combined forepaw touches. As impairment of the contralateral (i.e., right) forepaw was expected upon 6-OHDA lesion, a higher ratio of left to the sum of all touches indicates a more severe lesion. A ratio of 1.0 indicates that the mice only used the left forelimb (ipsilateral to the lesion) for the entire test session. Mice with a forepaw-touch ratio of less than 0.6 were considered poorly lesioned and were excluded from the study.

### *Ex vivo* electrophysiological recordings

Mice aged postnatal day 60–90 (4–6 weeks after AAV and 6-OHDA injections) were anesthetized with a ketamine-xylazine mixture and perfused transcardially with ice-cold artificial CSF (aCSF) containing the following (in mM): 125 NaCl, 2.5 KCl, 1.25 NaH_2_PO_4_, 2.0 CaCl_2_, 1.0 MgCl_2_, 25 NaHCO_3_, and 12.5 glucose, bubbled continuously with carbogen (95% O_2_ and 5% CO_2_). The brains were rapidly removed, glued to the stage of a vibrating microtome (Leica Instruments), and immersed in ice-cold aCSF. Parasagittal slices containing the GPe were cut at a thickness of 240 µm and transferred to a holding chamber, where they were submerged in aCSF with 3.33 mM pyruvate and 0.07 mM L-glutathione at 37°C for 30 min, and brought to room temperature before recording. Slices were then transferred to a small volume (0.5 ml) Delrin recording chamber that was mounted on a fixed-stage, upright microscope (Olympus). As there is no clear demarcation between the GPe and the more ventral structures in *ex vivo* brain slices, only neurons in the dorsal two-thirds of the GPe were sampled for electrophysiological analyses. GPe neurons were visualized using differential interference contrast optics, illuminated at 735 nm (Thorlabs), and imaged with a 60× 1.0 NA water-immersion objective (Olympus) and a CCD camera (QImaging). Genetically-labeled GPe neurons were identified based on their somatic tdTomato fluorescence and examined with epifluorescence microscopy using a white (6,500 K) LED (Thorlabs) and an appropriate filter cube (Semrock). Targeting of ChR2-eYFP was assessed by inspecting eYFP fluorescence before each recording. Mice with viral spread beyond STN and zona incerta were excluded from further experimentation.

Recordings were made at room temperature (20–22 °C) with patch electrodes fabricated from capillary glass (Sutter Instruments) pulled on a Flaming-Brown puller (Sutter Instruments) and fire-polished with a microforge (Narishige) immediately before use. Pipette resistance was typically 2–4 MOhms. The internal solution for cell-attached and voltage-clamp recordings consisted of the following (in mM): 125 CsMeSO_3_, 10 Na_2_-phosphocreatine, 5 tetraethylammonium chloride, 5 QX-314 Cl, 5 HEPES-K, 5 EGTA-K, 2 Mg_2_ATP, 0.5 CaCl_2_, 0.5 Na_3_GTP, 0.2% (w/v) biocytin, with pH adjusted to 7.25–7.30 with CsOH. SR95531 (10 µM) and CGP55845A (1 µM) were included in the bath during recordings to block GABAergic transmission. For SPN recordings Alexa Fluor 594 hydrazide (10–20 µM, Thermo Fisher Scientific) was included in the pipette solution to confirm the identity by visualizing the presence of spine-dense dendrites. Somatic patch-clamp recordings were obtained with an amplifier (Molecular Devices). The signal for all voltage-clamp recordings was filtered at 1 kHz and digitized at 10 kHz with a digitizer (Molecular Devices). For voltage-clamp recordings, series resistance was measured but not compensated for. The data were discarded if series resistance increased by 20% during recordings. Stimulus generation and data acquisition were performed using pClamp (Molecular Devices).

To measure the spontaneous and driven firing of GPe neurons, the cell-attached configuration was used to prevent the disruption of intracellular milieu. The average spontaneous firing was measured after two minutes of stable firing. To measure the driven firing of GPe neurons, input from the STN was optogenetically stimulated with 2 ms pulses at 10 Hz for 2 s. The average firing rate during the two-second stimulation period was taken as the driven firing rates of the respective cells. Whole-cell voltage-clamp recordings were used to measure excitatory postsynaptic currents (EPSCs). GPe neurons were voltage-clamped at –50 mV; terminals from excitatory inputs (from STN, PPN, and PF Thal) synapsed within the GPe were optogenetically stimulated for 2 ms. For *ex vivo* optogenetic experiments, blue excitation wavelength (peak, ∼450 nm) from two daylight (6,500 K) LEDs (Thorlabs) was delivered to the tissue slice from both a 60× water immersion objective and a 0.9 NA air condenser with the aid of 520 nm dichroic beamsplitters (Semrock). Light delivery was made at the site of electrophysiological recordings with a field of illumination of ∼500 µm in diameter. To measure STN input without the topographical biasing of the STN axons in the GPe, recordings were made from neighboring tdTomato^+^ and tdTomato^−^ neurons (less than 150 µm apart) in both PV-L-tdTom and Npas1-Cre-tdTom mice. To study STN input with CreOn expression of ChR2, EPSC recordings were made from tdTomato^+^ and tdTomato^−^ GPe neurons in PV-L-tdTom mice. Corticostriatal EPSCs were evoked with electrical stimulation using parallel bipolar tungsten electrodes (Frederick Haer) placed in the cortex. To measure AMPA and NMDA receptor-mediated currents, the holding potential of recorded neurons was alternated between –80 mV and +40 mV. AMPA receptor-dependent currents were measured from the peak amplitude of EPSCs at –80 mV. NMDA receptor-dependent currents were measured at 50 ms after the EPSC peak at +40 mV. AMPA-NMDA ratio was calculated by dividing the AMPA current to NMDA current.

### Histology

For histological verification of injection and implantation sites in mice used for *in vivo* behavioral experiments, mice were anesthetized deeply with a ketamine-xylazine mixture and perfused transcardially first with 0.01 M phosphate-buffered saline followed by a fixative containing 4% paraformaldehyde (PFA, pH 7.4). Brain tissue was then postfixed in the same fixative for 2 h at 4°C. Tissue blocks containing the GPe were sectioned sagittally using a vibrating microtome (Leica Instruments) at a thickness of 60 µm. Sections were then washed, mounted, and coverslipped. Standard immunohistological procedures (Abecassis et al., 2020) were performed in a subset of brain tissues to validate the cellular specificity of transgene expression. Sections were imaged under an epifluorescence microscope (Keyence), and each section was visually inspected. Graphical representations were created using a thresholding based protocol in Fiji (http://fiji.sc/Fiji) (Schindelin et al., 2012), where regions of interest were selected to show transgene expression in the primary injection sites. For histological verification of AAV_retro_-mediated recombination and the quantification of STN fibers, images were taken on a laser-scanning confocal microscope with a 10× 0.45 numerical aperture (NA) air objective and a 60× 1.35 NA oil-immersion objective (Olympus).

To quantify STN axonal fibers in the GPe, identical histological procedures were used. Sections were imaged on a laser-scanning confocal microscope with a 10× 0.45 NA air objective (Olympus). Values for ‘area’, ‘area fraction,’ and ‘integrated density’ were quantified using Fiji. Specifically, area represents the user-defined area of the GPe in a sagittal section, and area fraction represents the fraction of area that contains pixels with values above the set threshold; in this case, pixels with eYFP-labeled STN fibers. Integrated density represents the sum of all pixels within the selected area, i.e., the sum of absolute pixel values, which was used as a measure of the density of STN fibers within the GPe. To quantify putative STN terminals within the GPe, standard immunohistological procedures were performed. Floating tissue sections were reacted with antibodies for GFP (1:1,000; Abcam) and VGluT2 (1:1,000; NeuroMab). Images were taken on a laser-scanning confocal microscope with a 60× 1.35 NA oil-immersion objective and 2x digital zoom. Three non-overlapping fields (106.7 x 106.7 µm^2^) were imaged from each section; 5-μm serial optical sections (z-stacks, at 1 μm intervals) were acquired. Puncta were deemed *bona fide* STN-GPe synaptic boutons only if they showed spatial overlap of eYFP and VGluT2 signals across all three orthogonal planes. GPe sections from three different equally-spaced (400 µm) lateromedial levels were sampled (Hernandez et al., 2015; Abecassis et al., 2020) from the left hemisphere.

Brain tissue used for *ex vivo* electrophysiological recordings were stored in 4% paraformaldehyde for 24 hours. PFA-fixed tissues were mounted on microscope slides, coverslipped with Prolong Gold antifade mounting medium (Thermo Fisher Scientific), and imaged under an epifluorescence microscope (Keyence) to inspect the fluorescent signal from the site of viral injection.

### Experimental design and statistical analyses

Statistical analyses were performed with MATLAB (Mathworks) and Prism (GraphPad). Custom analysis codes are available on GitHub (https://github.com/chanlab). The sample size (n value) shown represents the number of mice for *in vivo* behavior experiments and the number of cells for *ex vivo* electrophysiological experiments. Unless noted otherwise, data are presented as median values ± median absolute deviations (MADs) (Leys et al., 2013) as measures of central tendency and statistical dispersion, respectively. Box plots are used for graphical representation of the distribution of values (Krzywinski and Altman, 2014; Streit and Gehlenborg, 2014; Nuzzo, 2016). In a box plot, the central line represents the median, the box edges represent the interquartile ranges, and whiskers represent 10–90^th^ percentiles. No statistical method was used to predetermine sample sizes.

Comparisons for unpaired samples were performed using a Mann–Whitney U test. Comparisons for paired samples were performed using the Wilcoxon signed-rank test. The Fisher’s exact test was used to determine if there were nonrandom associations in two nominal variables with a threshold (α) of 0.05 for significance. Statistical significance was set at P < 0.05. Unless < 0.0001 or > 0.99, exact *P* values (two-tailed) are reported. No multiple testing corrections were considered.

## Results

We sought to dissect the relationship between GPe neuron activity and motor output with cell type-specific strategies. To confer transgene expression specificity in PV^+^ neurons and Npas1^+^ neurons, we used Cre recombinase-inducible expression (CreOn) adeno-associated viruses (AAVs) in conjunction with PV-Cre and Npas1-Cre-tdTom mouse lines, respectively. *In vivo* optogenetics, which provides a relatively high temporal resolution, was used to interrogate the roles that PV^+^ neurons and Npas1^+^ neurons play in regulating full-body movements—locomotion. Locomotor activity of mice in an open-field arena across individual trials was analyzed. We focused on three defined time windows relative to light delivery: the ‘pre-period’ (pre, −10 to 0 s), the ‘light-period’ (light, 0 to 10 s), and ‘post-period’ (post, 10 to 20 s).

### PV^+^ neurons promote movement

Stimulation of PV^+^ neurons *in vivo* using ChR2 (an excitatory opsin) (Boyden et al., 2005) (**Figure 1a**) induced an increase in motor output, as measured by the duration of movement periods (not shown) and locomotion speed (**Figure 1b–c**, **Movie 1**). Both patterned and sustained activation of ChR2 were effective in promoting speed (patterned: +0.52 ± 0.31 fold, n = 11 mice, P = 0.0049; sustained: +0.79 ± 0.28 fold, n = 12 mice, P = 0.00049). The motor response induced by the two protocols were not statistically different (P = 0.094). The motor effect was readily reversed following the cessation of light delivery, as the speed during the post-period was not different from that during the pre-period. The motor response was not the result of non-specific effects of light delivery, such as heat artifact (Owen et al., 2019), as PV-Cre mice transduced with a control virus (eYFP only) did not display any motor effects with light delivery (patterned: –0.035 ± 0.25 fold, n = 13 mice, P = 0.84; sustained: –0.14 ± 0.20 fold, n = 13 mice, P = 0.17) (**Figure 1b**, **2a**, **Movie 2**). Similarly, sham-injected PV-Cre mice did not show any motor responses associated with light delivery (patterned: +0.11 ± 0.20 fold, n = 11 mice, P = 0.28; sustained: –0.16 ± 0.31 fold, n = 9 mice, P = 0.91).

**Figure 1.**
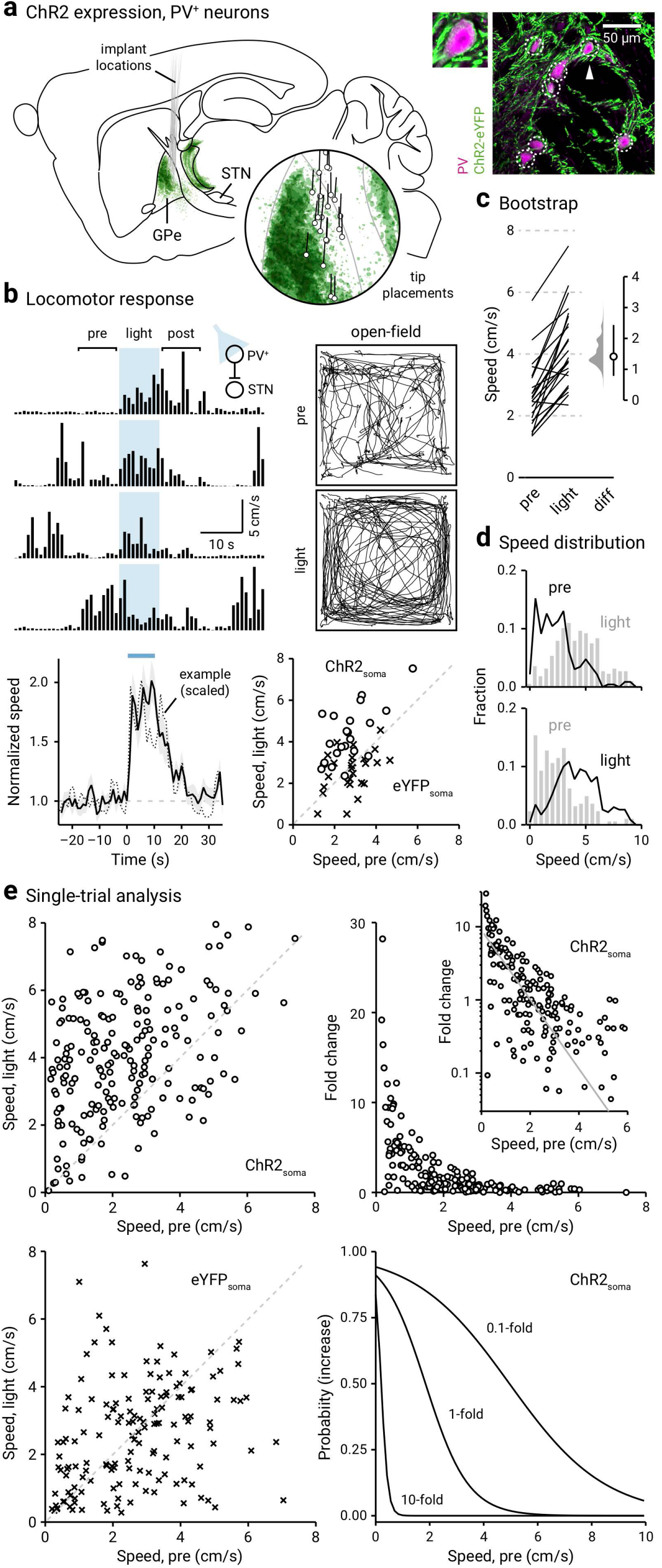
**Optogenetic stimulation of PV^+^ neurons promotes locomotion**. **a.** Left: Representation of ChR2-eYFP expression patterns and fiber optic implant locations in PV-Cre mice (n = 12). Inset: Representation of optical fiber tip placements in relation to the GPe and neighboring structures. Right: Confocal micrographs showing the cellular specificity of ChR2-eYFP expression in the GPe. For clarity, a magnified example (arrowhead) is shown. Abbreviations: GPe, external globus pallidus; STN, subthalamic nucleus. **b.** Top, left: A representative example of the locomotor activity of a single mouse across four trials. The schematic diagram shows the site of light delivery. Blue shaded area indicates the duration (10 s) of light delivery. Top, right: Movement tracks corresponding to the ‘pre-period’ (top) and ‘light-period’ (bottom). Six representative mice (ten trials from each) are presented. Bottom, left: A plot showing the relationship between normalized speed and time. Blue bar indicates the duration (10 s) of light delivery. The dotted horizontal line indicates the baseline locomotor activity level. Black solid trace is the population mean calculated from all mice; shading indicates the standard error of the mean. Black dotted trace is a representative example from a single mouse; data were scaled to facilitate comparison. Bottom, right: Speed during ‘light-period’ against speed during ‘pre-period’ is plotted. Data from PV-Cre mice expressing ChR2-eYFP (circles) and eYFP (crosses) are displayed. Each marker represents a mouse. The diagonal line indicates unity (i.e., x = y). Data points for ChR2-eYFP are systematically shifted upwards relative to unity. **c.** Slopegraph showing the response of each mouse upon optogenetic stimulation of PV^+^ neurons. The slope of each line shows the magnitude and direction of change. Median difference is plotted on a floating axis. The smoothed density represents bootstrap resampled data. The median difference is indicated as a circle and the 95% confidence interval is indicated by the length of vertical lines. **d.** Speed distributions of mice during ‘pre-period’ and ‘light-period’ are shown. **e.** Top, left: A scatter plot showing the pairwise relationship between speed during ‘light-period’ and speed during ‘pre-period’ from PV-Cre mice expressing ChR2-eYFP. Each marker is a trial. The diagonal line indicates unity. Top right: Fold change in speed with light delivery against speed during ‘pre-period’ from PV-Cre mice expressing ChR2-eYFP. Each marker is a trial. Inset: Same data displayed with fold increase on a log scale. Grayline is a monoexponential fit of the data. Bottom, left: A plot showing the speed during ‘light-period’ versus speed during ‘pre-period’. Data from PV-Cre mice expressing eYFP are displayed. Each marker is a trial. The diagonal line indicates unity. Bottom, right: Probability of increase against speed during ‘pre-period’. Logistic regression curves fitted to data for 0.1-fold, 1-fold, and 10-fold increases are displayed.

By examining the statistics of spontaneous (pre-period) and induced locomotion (light-period), it became clear that optogenetic stimulation induced a rightward shift in the speed distribution (**Figure 1d**). Furthermore, mice also showed a decrease in time spent immobile (i.e., horizontal displacement ≤ 1 cm/s) upon optogenetic stimulation of PV^+^ neurons (pre-period: 34 ± 21%, light-period: 11 ± 9.5%, n = 230 trials, P < 0.0001). Though mice spend a larger fraction of time ambulating at higher velocities during induced locomotion, the activity level spanned the same range as observed during spontaneous locomotion. These salient features of the relationship between the activity during pre-period and light-period are not always captured at single-animal analyses. As demonstrated by the example in **Figure 1b**, strong motor effects were observed in trials only when low ambulatory levels occurred immediately preceding the optogenetic stimulus. To gain deeper insights into how spontaneous locomotor activity level shaped the size of the motor response, we analyzed data from individual trials. As shown in **Figure 1e**, the speed in the light-period was distinct from their pre-period level; most of the data points are above unity. On the contrary, eYFP controls did not display any motor effects associated with light delivery (bottom left); all data points are distributed evenly along the unity. An important feature in these data is the strong negative relationship between pre-stimulus speed and the fold increase in locomotion. As illustrated in **Figure 1e**, this relationship can be described by a monoexponential function. Furthermore, logistic regression predicted the motor effect when pre-stimulus speed was considered as a covariate (**Figure 1e**, **Table 4**). *Post hoc* histological examination confirmed that the behavioral effects were due to specific targeting of the GPe (**Figure 1a**). As viral spread to the thalamic reticular nucleus (TRN) was apparent in some cases, we examined whether our results were confounded. Using near-identical procedures, we found that selective ChR2 expression in the TRN did not yield consistent motor effects (patterned: –0.12 ± 0.22 fold, n = 11 mice, P = 0.90; sustained: –0.17 ± 0.21 fold, n = 11 mice, P = 0.41) (**Figure 2a & b**).

**Figure 2.**
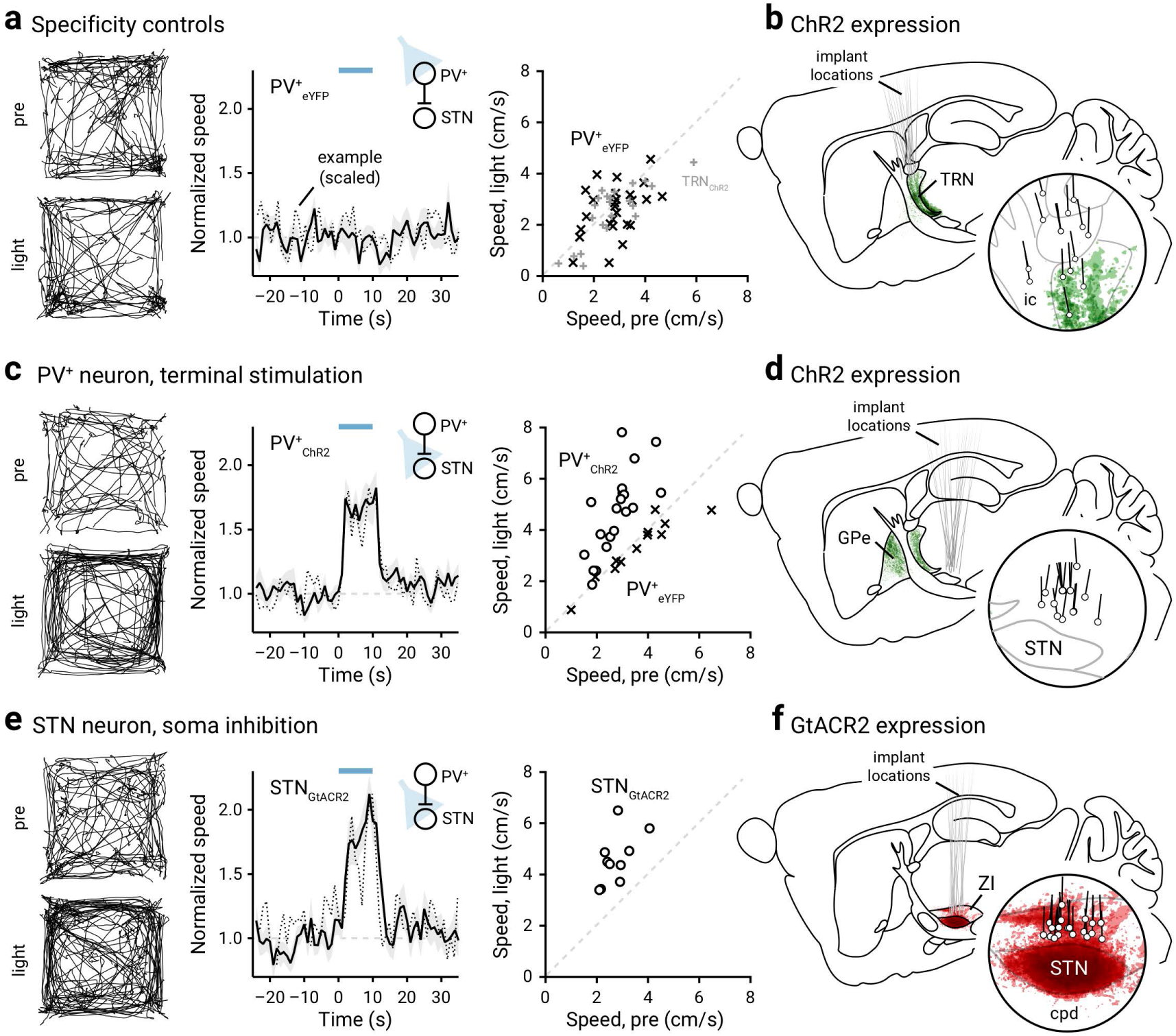
Inhibition of STN promotes locomotion. **a.** Left: Movement tracks corresponding to the ‘pre-period’ (top) and ‘light-period’ (bottom). Six representative mice (ten trials from each) are presented. Middle: A plot showing the relationship between normalized speed and time. Blue bar indicates the duration (10 s) of light delivery. The dotted horizontal line indicates the baseline locomotor activity level. Black solid trace is the population mean calculated from all mice; shading indicates the standard error of the mean. Black dotted trace is a representative example from a single mouse; data were scaled to facilitate comparison. The same presentation scheme is used in **c** and **e**. Data from eYFP-expressing PV-Cre mice are shown; light was delivered to the GPe. Data from the same mice (crosses) are presented on the right in a scatterplot, which shows the speed during light delivery versus speed during ‘pre-period’. Mice with ChR2-expressed in thalamic reticular nucleus (TRN) neurons are used as additional controls (gray pluses). **b.** Representation of targeted ChR2-eYFP expression patterns and fiber optic implant locations in PV-Cre mice (n = 11). The TRN was targeted in these experiments. **c.** Left, middle: Data from optogenetic stimulation of PV^+^ neuron terminals in the STN are shown. Right: eYFP-expressing mice (in PV^+^ neurons) were used as controls (crosses). **d.** Representation of ChR2-eYFP expression patterns and fiber optic implant locations in PV-Cre mice (n = 10). ChR2-expressing PV^+^ neuron terminals in the STN were targeted. **e.** Data from optogenetic inhibition of STN neurons with GtACR2 are shown. **f.** Representation of GtACR2-FusionRed expression patterns and fiber optic implant locations in C57BL/6J mice (n = 10). The STN was targeted in these experiments. Abbreviations: cpd, cerebral peduncle; GPe, external globus pallidus; ic, internal capsule; STN, subthalamic nucleus; TRN, thalamic reticular nucleus; ZI, zona incerta.

**Table 4.** Summary of logistic regression models.

| Strain, manipulation | Fold changes | Odds ratio (OR) | P value | 95% confidence interval for ORs |  |
| --- | --- | --- | --- | --- | --- |
|  |  |  |  | Lower bound | Upper bound |
| PV-Cre ChR2 | 0.1-fold increase | 0.569 | **< 0.001** | 0.465 | 0.696 |
|  | 1-fold increase | 0.283 | **< 0.001** | 0.201 | 0.398 |
|  | 10-fold increase | 3.319e-4 | **0.0093** | 0.000 | 0.138 |
| PV ChR2, 6-OHDA | 0.1-fold increase | 0.673 | **0.0012** | 0.530 | 0.856 |
|  | 1-fold increase | 0.388 | **< 0.001** | 0.278 | 0.541 |
|  | 10-fold increase | 9.736e-7 | **0.0012** | 0.000 | 0.004 |
| Npas1-Cre-tdTom ChR2 | 0.1-fold decrease | 1.589 | **< 0.001** | 1.288 | 1.960 |
|  | 0.5-fold decrease | 1.589 | **< 0.001** | 1.288 | 1.960 |
|  | 0.8-fold decrease | 1.227 | 0.16 | 0.923 | 1.631 |

The bolded P values are significant (i.e. < 0.05).

PV^+^ neurons send inhibitory projections primarily to the STN and SNr (Bevan et al., 1998; Sato et al., 2000a; Kita, 2007; Mastro et al., 2014; Hernandez et al., 2015; Fujiyama et al., 2016; Saunders et al., 2016; Oh et al., 2017; Abecassis et al., 2020). To confirm that the movement-promoting effects of PV^+^ neuron stimulation were mediated through inhibiting these downstream targets, we optogenetically stimulated PV^+^ neuron axon terminals in the STN (**Figure 2c & d**). As expected, stimulation of PV^+^ axon terminals in the STN resulted in an increase in the speed of movement (patterned: +0.41 ± 0.16 fold, n = 10 mice, P = 0.0020; sustained: +0.84 ± 0.11 fold, n = 10 mice, P = 0.0020). Light delivery at the PV^+^ axon terminals in the STN of PV-Cre mice transduced with a control virus (eYFP only) did not produce any motor effects with light delivery (patterned: –0.0052 ± 0.10 fold, n = 6 mice, P > 0.99; sustained: –0.081 ± 0.027 fold, n = 6 mice, P = 0.31) (**Figure 2c**). To exclude action potential propagation as a confound, we activated virally-expressed GtACR2 (an inhibitory opsin) (Govorunova et al., 2015; Mahn et al., 2018) in STN neurons to mimic the inhibitory action of the GPe-STN input. As expected, we observed an increase in movement (+0.60 ± 0.16 fold, n = 10 mice, P = 0.0020) (**Figure 2e, 2f**, **Movie 3**). In sum, our data collectively argue that activity of PV^+^ neurons promoted locomotion via the inhibition of their primary downstream targets.

### Npas1^+^ neurons suppress movement

In contrast with the motor effect of PV^+^ neurons, *in vivo* optogenetic stimulation of Npas1^+^ neurons induced a decrease in the duration of movement periods (not shown) and locomotion speed (patterned: –0.41 ± 0.10 fold, n = 14 mice, P = 0.000031; sustained: –0.37 ± 0.089 fold, n = 16 mice, P = 0.00024) (**Figure 3a–c**, **Movie 4**). The effect of optogenetic stimulation of Npas1^+^ neurons with both a sustained light pulse and a 20 Hz-train were not different (P = 0.24) in suppressing speed as the measure of motor output. This observation is consistent with our previous findings with chemogenetic stimulation of Npas1^+^ neurons (Glajch et al., 2016). Importantly, our observations provide a causal demonstration of the proposed role of Npas1^+^ neurons in movement suppression (Mallet et al., 2016). The optogenetically-induced motor suppression was readily reversible, as the speed during the post-period was not different from that during the pre-period (**Figure 3b**).

**Figure 3.**
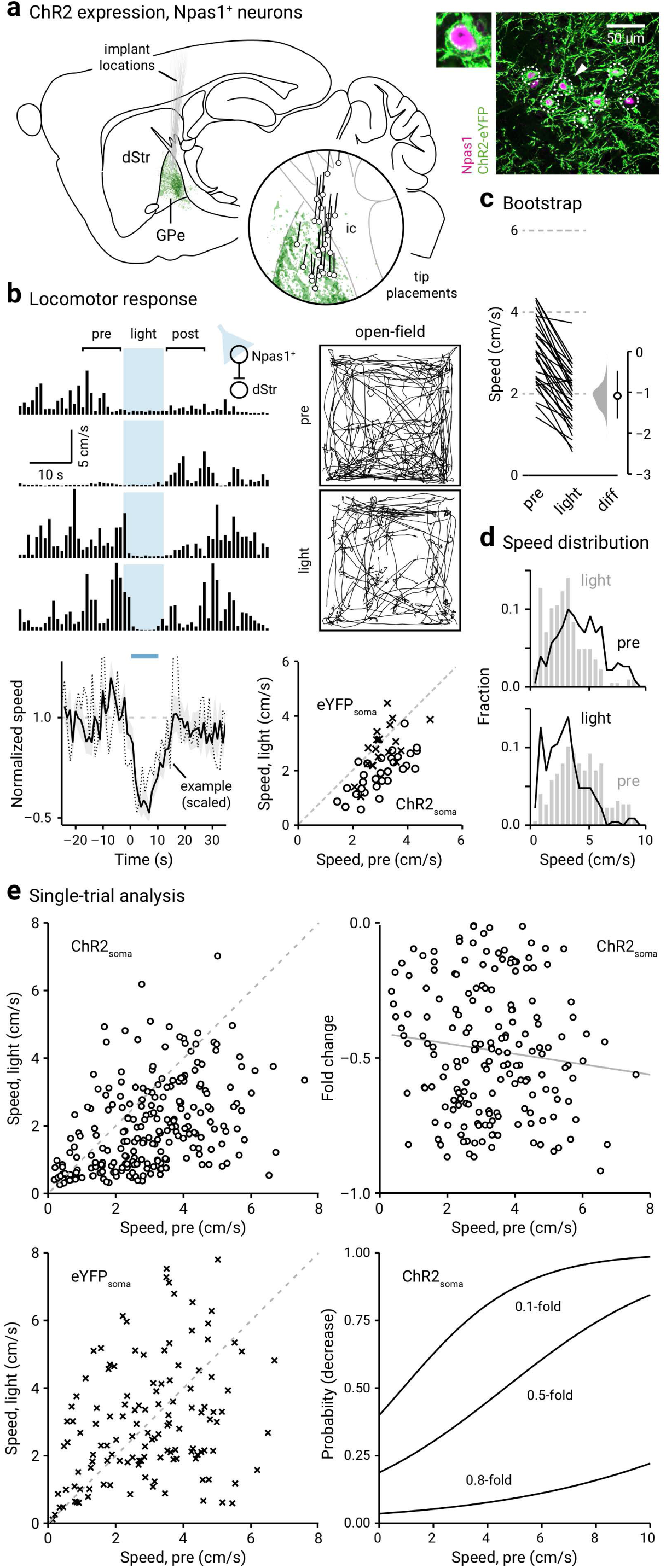
Optogenetic stimulation of Npas1^+^ neurons suppresses locomotion. **a.** Left: Representation of ChR2-eYFP expression patterns and fiber optic implant locations in Npas1-Cre-tdTom mice (n = 14). Inset: Representation of optical fiber tip placements in relation to the GPe and neighboring structures. Right: Confocal micrographs showing the cellular specificity of ChR2-eYFP expression in the GPe. For clarity, a magnified example (arrowhead) is shown. Abbreviations: dStr, dorsal striatum; GPe, external globus pallidus; ic, internal capsule. **b.** Top, left: A representative example of the locomotor activity of a single mouse across four trials. The schematic diagram shows the site of light delivery. Blue shaded area indicates the duration (10 s) of light delivery. Top, right: Movement tracks corresponding to the ‘pre-period’ (top) and ‘light-period’ (bottom). Six representative mice (ten trials from each) are presented. Bottom, left: A plot showing the relationship between normalized speed and time. Blue bar indicates the duration (10 s) of light delivery. The dotted horizontal line indicates the baseline locomotor activity level. Black solid trace is the population mean calculated from all mice; shading indicates the standard error of the mean. Black dotted trace is a representative example from a single mouse; data were scaled to facilitate comparison. Bottom, right: Speed during ‘light-period’ against speed during ‘pre-period’ is plotted. Data from Npas1-Cre-tdTom mice expressing ChR2-eYFP (circles) and eYFP (crosses) are displayed. Each marker represents a mouse. The diagonal line indicates unity (i.e., x = y). Data points for ChR2-eYFP are systematically shifted downwards relative to unity. **c.** Slopegraph showing the response of each mouse upon optogenetic stimulation of Npas1^+^ neurons. The slope of each line shows the magnitude and direction of change. Median difference is plotted on a floating axis. The smoothed density represents bootstrap resampled data. The median difference is indicated as a circle and the 95% confidence interval is indicated by the length of vertical lines. **d.** Speed distributions of mice during ‘pre-period’ and ‘light-period’ are shown. **e.** Top, left: A scatter plot showing the pairwise relationship between speed during ‘light-period’ and speed during ‘pre-period’ from Npas1-Cre-tdTom mice expressing ChR2-eYFP. Each marker is a trial. The diagonal line indicates unity. Top, right: Fold change in speed with light delivery against speed during ‘pre-period’ from Npas1-Cre-tdTom mice expressing ChR2-eYFP. Each marker is a trial. Grayline is a linear fit of the data. Bottom left: A plot showing the speed during ‘light-period’ versus speed during ‘pre-period’. Data from Npas1-Cre-tdTom mice expression eYFP are displayed. Each marker is a trial. The diagonal line indicates unity. Bottom, right: Probability of decrease against speed during ‘pre-period’. Logistic regression curves fitted to data for 0.1-fold, 0.5-fold, and 0.8-fold decreases are displayed.

By visualizing the speed distributions during the pre-period and light-period, it is apparent that in addition to a leftward shift in the speed distribution (**Figure 3d**), mice also showed an increase in time spent immobile upon optogenetic stimulation of Npas1^+^ neurons (pre-period: 26 ± 16%, light-period: 46 ± 24%, n = 240 trials, P = 0.0020). Similar to that observed in PV^+^ neurons, single-trial data revealed that the effect size was a function of the level of spontaneous activity (i.e., during pre-period) immediately preceding the light-period. Strong motor suppression was associated with a high level of spontaneous movement preceding the light-period. This relationship can be described by a linear function (**Figure 3e**). The magnitude of motor suppression was predicted by the level of spontaneous movement with logistic regression (**Figure 3e**, **Table 4**). On the other hand, Npas1-Cre-tdTom mice transduced with a control AAV (eYFP only) did not exhibit any changes in locomotor activity with light delivery (patterned: +0.06 ± 0.19 fold, n = 11 mice, P = 0.83; sustained: –0.13 ± 0.15 fold, n = 10 mice, P = 0.13) (**Figure 3b**, **4a**, **Movie 5**). Similarly, sham-injected Npas1-Cre-tdTom mice also did not show any motor responses associated with light delivery (patterned: +0.020 ± 0.081 fold, n = 6 mice, P > 0.99; sustained: +0.19 ± 0.36 fold, n = 6 mice, P > 0.99).

Next, we wanted to ask if Npas1^+^ neurons can bidirectionally regulate motor output. *In vivo* optogenetic inhibition of Npas1^+^ neurons using GtACR2 induced an increase in movement (+0.34 ± 0.34 fold, n = 9 mice, P = 0.020) (**Figure 4b**), arguing that Npas1^+^ neurons were exerting an ongoing inhibition of their downstream targets *in vivo*, during spontaneous locomotion. Npas1^+^ neurons primarily project to the dStr (Kita et al., 1999; Abdi et al., 2015; Dodson et al., 2015; Hernandez et al., 2015; Saunders et al., 2016; Abecassis et al., 2020). As the dStr is responsible for motor behavior (Kravitz et al., 2010; Durieux et al., 2012; Freeze et al., 2013), we stimulated Npas1^+^ terminals within the dStr to determine whether the movement-suppressing effects of stimulation of Npas1^+^ neurons are mediated through this downstream projection. Similar to the effects observed with somatic stimulation of Npas1^+^ neurons, optogenetic stimulation of their terminals in the dStr led to a reduction in movement (patterned: –0.29 ± 0.11 fold, n = 12 mice, P = 0.00049; sustained: –0.19 ± 0.14 fold, n = 12 mice, P = 0.034) (**Figure 4c**). In contrast, light delivery at the Npas1^+^ neuron terminals in the dStr of Npas1-Cre-tdTom mice transduced with a control AAV (eYFP only) did not result in any motor effects (patterned: +0.058 ± 0.17 fold, n = 11 mice, P = 0.77; sustained: +0.065 ± 0.16 fold, n = 11 mice, P = 0.32).

**Figure 4.**
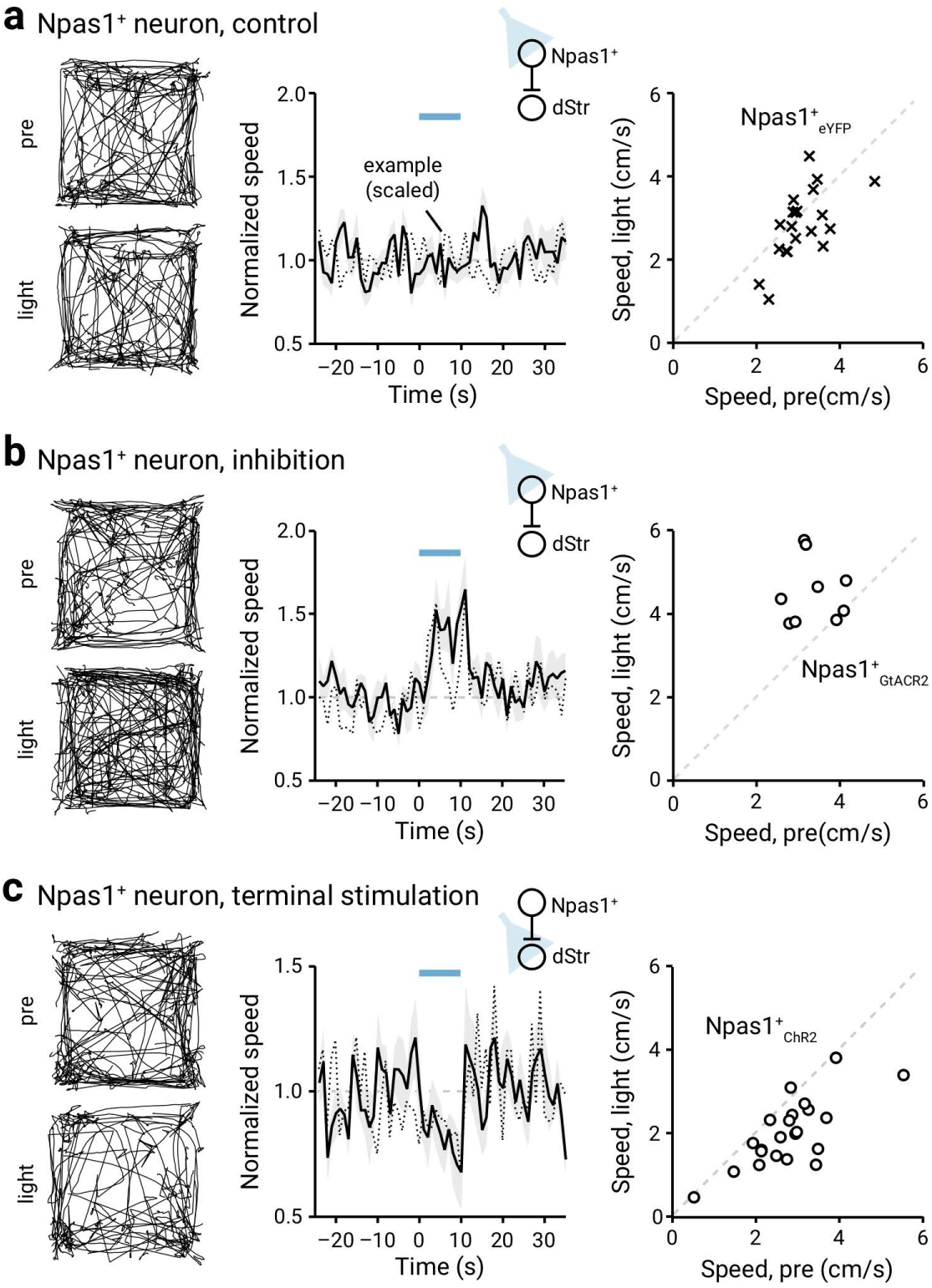
Ongoing Npas1^+^ neuron activity suppresses locomotion. **a.** Left: Movement tracks corresponding to the ‘pre-period’ (top) and ‘light-period’ (bottom). Six representative mice (ten trials from each) are presented. Middle: A plot showing the relationship between normalized speed and time. Blue bar indicates the duration (10 s) of light delivery. The dotted horizontal line indicates the baseline locomotor activity level. Black solid trace is the population mean calculated from all mice; shading indicates the standard error of the mean. Black dotted trace is a representative example from a single mouse; data were scaled to facilitate comparison. The same presentation scheme is used in **b** and **c**. Data from eYFP-expressing Npas1-Cre-tdTom mice are shown; light was delivered to the GPe. **b.** Data from GtACR2-expressing Npas1-Cre-tdTom mice are shown; light was delivered to the GPe. **c.** Data from optogenetic stimulation of Npas1^+^ neuron terminals in the dorsal striatum (dStr) are shown.

To confirm that the motor effect induced by optogenetic manipulation of GPe output were not a result of settings unique to our experimental setup, we optogenetically stimulated direct and indirect pathway striatal projection neurons (dSPNs and iSPNs, respectively) in the dStr. This resulted in a canonical increase (+1.73 ± 0.21 fold, n = 8 mice, P = 0.0078) and decrease (–0.61 ± 0.077 fold, n = 8 mice, P = 0.0078) in movement, respectively, as demonstrated previously (Kravitz et al., 2010). To summarize, we conclude that ongoing Npas1^+^ neuron activity modulates motor output — stimulation of Npas1^+^ neurons suppressed locomotion.

### The STN preferentially targets PV^+^ neurons

Our *in vivo* optogenetic interrogation showed that PV^+^ neurons are movement-promoting and Npas1^+^ neurons are movement-suppressing. However, the excitatory inputs that naturally drive the activity of these GPe neurons have not been fully characterized. Both anatomical and physiological studies show that the principal glutamatergic input to the GPe arises from the STN (Carpenter et al., 1981a; Carpenter et al., 1981b; Kita and Kitai, 1987; Smith et al., 1990a; Smith et al., 1990b; Smith et al., 1998; Kita et al., 2004; Koshimizu et al., 2013). However, it is not known whether the STN selectively targets particular GPe neuron subpopulations. Computational models suggest that the STN targets a select subset of Npas1^+^ neurons (Bogacz et al., 2016); however, this hypothesis was yet to be empirically tested.

To study the properties of the STN-GPe input, we performed whole-cell, voltage-clamp recordings from genetically-identified PV^+^ neurons and Npas1^+^ neurons in an acute brain slice preparation. To allow the stimulation of the STN input, an optogenetic approach was employed. Injection of the STN with an AAV that expressed ChR2-eYFP constitutively led to robust ChR2-eYFP expression in STN neurons. This gave rise to eYFP-labeled axonal processes terminating throughout the GPe. The VGluT2-immunoreactive puncta on these axons suggest that they are terminating within the GPe and have the capacity to release glutamate (**Figure 5a–b**). The properties of STN inputs to GPe were examined in the presence of GABAergic antagonists (10 µM SR95531, 1 µM CGP55845). Optogenetic stimulation of the STN input reliably evoked excitatory postsynaptic currents (EPSCs) in all neurons tested. Notably, EPSC amplitudes, as a measure of the STN input strength, were roughly five times larger in PV^+^ neurons compared to those in Npas1^+^ neurons (PV^+^ = 674.01 ± 174.47 pA, n = 34 neurons; Npas1^+^ = 128.30 ± 63.05 pA, n = 41 neurons; P < 0.0001) (**Figure 5d & e**). The EPSCs evoked with the constitutive ChR2 expression were not due to ectopic expression of ChR2 in the zona incerta (ZI), which is immediately adjacent (dorsocaudal) to the STN, as selectively-targeted injection and optogenetic stimulation of the ZI input did not produce large EPSCs in either PV^+^ neurons or Npas1^+^ neurons (PV^+^: 33.93 ± 27.22 pA, n = 9 neurons; Npas1^+^: 33.08 ± 20.39 pA, n = 9 neurons; P = 0.26) (**Figure 5c–d**).

**Figure 5.**
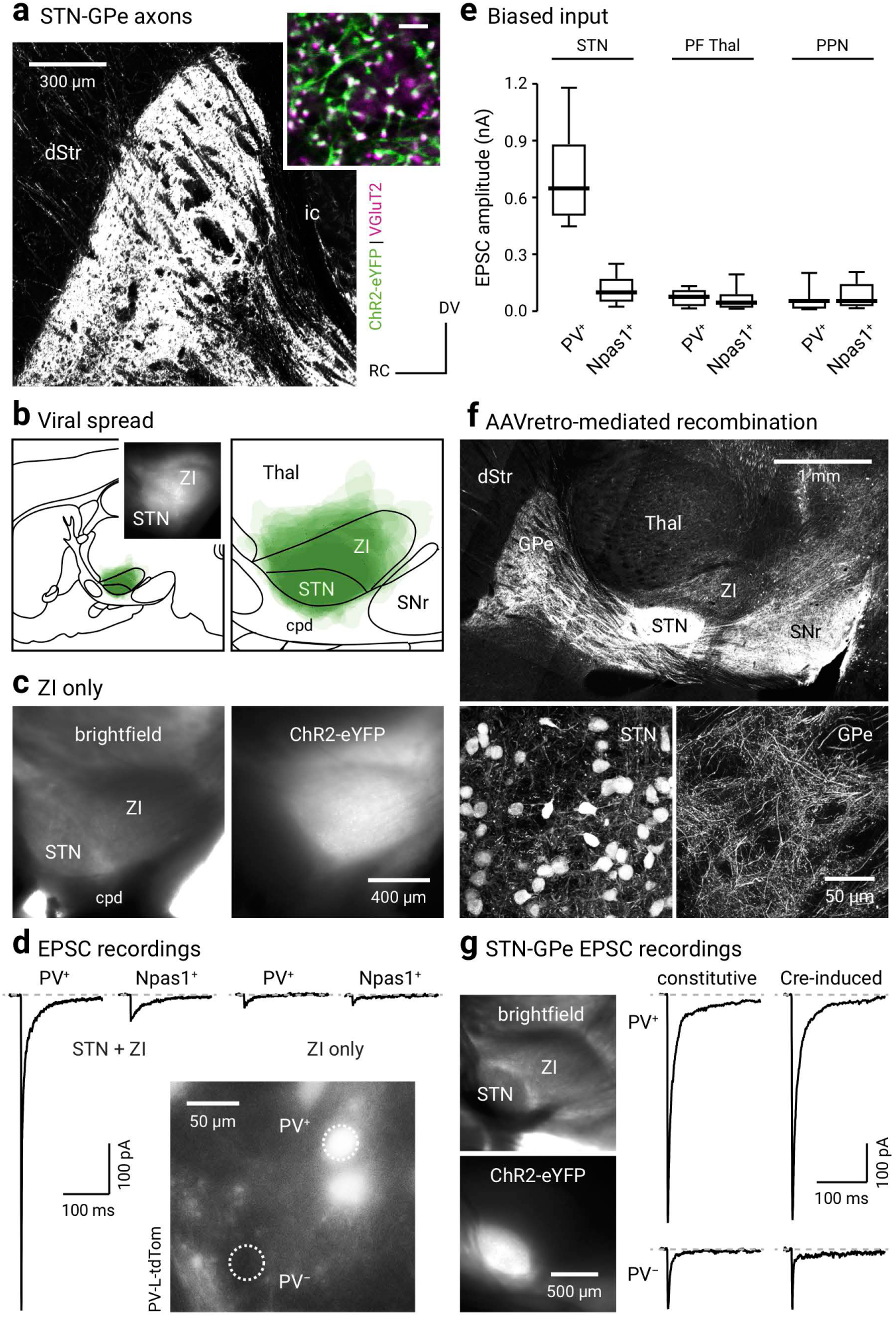
STN input is biased toward PV^+^ neurons. **a.** A confocal micrograph of a sagittal brain section showing eYFP-labeled subthalamic nucleus (STN) axons in the GPe (in a mouse injected with a ChR2-eYFP AAV in the STN). Inset: High magnification confocal micrograph showing the spatial relationship between eYFP-labeled STN axons (green) and VGluT2 (magenta) labeling. Scale bar = 5 µm. **b.** Left: A schematic showing viral spread overlaid for each subject (n = 9) used for *ex vivo* experiments. Inset: A representative epifluorescent image of a parasagittal acute brain slice showing the expression of ChR2-eYFP in the STN and the neighboring areas. Right: A magnified view of transduced areas is shown. **c.** Left: A brightfield image of a parasagittal acute brain slice showing STN and ZI. Right: eYFP signal from the same slice showing transduction only in the ZI but not the STN. **d.** Excitatory postsynaptic currents (EPSCs) recorded in PV^+^ neurons and Npas1^+^ neurons in mice that had STN + ZI and ZI only transductions. Inset: Epifluorescence image from *ex vivo* tissue showing the GPe of a PV-L-tdTom mouse with tdTomato+ (PV^+^) and tdTomato^−^ (PV^−^) neurons within the same field. **e.** Box plots summarizing the amplitude of EPSCs recorded in PV^+^ neurons and Npas1^+^ neurons with optogenetic stimulation of terminals from the STN, the parafascicular nucleus of the thalamus (PF Thal), or the pedunculopontine nucleus (PPN). **f.** Top: A montage of confocal micrographs from a sagittal brain section in a mouse injected with AAV_retro_-Cre in the SNr, along with Cre recombinase-inducible expression (CreOn) ChR2-eYFP-expressing AAVs in the STN. These images show that eYFP-labeled STN axons are arborized throughout the GPe and SNr. Bottom: High magnification images showing eYFP-labeled neurons in the STN (left) and their axons in the GPe (right). **g.** Left: CreOn expression of ChR2-eYFP in the STN (top), where eYFP-labeling (bottom) is localized within the STN. Right: Representative EPSC recordings from voltage-clamped PV^+^ neurons (top) and PV^−^ neurons (bottom) in mice with constitutive (left) or CreOn (right) expression of ChR2-eYFP in the STN. Abbreviations: cpd, cerebral peduncle; dStr, dorsal striatum; GPe, external globus pallidus; ic, internal capsule; STN, subthalamic nucleus; SNr, substantia nigra pars reticulata; Thal, thalamus; ZI, zona incerta.

As the methodology employed did not rely on an STN-specific driver or promoter, a Cre-lox strategy was also used to verify the specificity of the involvement of the STN in our analysis. To efficiently deliver Cre to STN neurons, we injected AAV_retro_-Cre (Tervo et al., 2016) in the SNr, which is the principal downstream target of the STN (Parent and Smith, 1987; Smith et al., 1998). This Cre-delivery strategy yielded robust recombination events in STN neurons, as depicted by tdTomato expression in the Cre reporter line (LSL-tdTomato) (**Figure 5f**). PV-tdTom mice were then used in subsequent experiments to allow for the identification of PV^+^ neurons (Abecassis et al., 2020). By using the same retrograde Cre-delivery strategy in conjunction with CreOn ChR2-expressing AAVs (injected into the STN), EPSCs were reliably evoked when STN input was optogenetically stimulated. Consistent with our observations using constitutive expression of ChR2 in the STN, EPSCs evoked with the AAV_retro_-Cre-mediated strategy were larger in PV^+^ neurons than those in PV^−^ neurons (PV^+^: 578.92 ± 49.01 pA, n = 6 neurons; PV^−^: 155.64 ± 52.61 pA, n = 5 neurons; P = 0.0043). Moreover, EPSCs measured in PV^+^ neurons and PV^−^ neurons in mice with Cre-mediated ChR2 expression in the STN were not different than those measured in PV^+^ neurons and Npas1^+^ neurons in mice with constitutive ChR2 expression in the STN (PV^+^ vs PV^+^: P = 0.12; PV^−^ vs Npas1^+^: P = 0.51) (**Figure 5g**). In summary, our experiments confirmed the validity of the general approach employed in this study.

In addition to the STN, the parafascicular nucleus of the thalamus (PF Thal) and pedunculopontine nucleus (PPN) are known to send glutamatergic projections to the GPe (Kincaid et al., 1991; Sadikot et al., 1992; Naito and Kita, 1994; Deschenes et al., 1996; Mouroux et al., 1997; Kita et al., 2004; Yasukawa et al., 2004). We sought to compare the properties of these inputs to the STN input using near-identical approaches (see Methods). In contrast to the STN input, PF Thal and PPN inputs did not produce detectable EPSCs in all GPe neurons. Fewer PV^+^ neurons (90.6%, 19 out of 21 neurons) and Npas1^+^ neurons (88.2%, 15 out of 17 neurons) responded to PF Thal input stimulation (PV^+^ = 76.64 ± 44.3 pA, n = 19 neurons; Npas1^+^ = 45.32 ± 26.9 pA, n = 15 neurons) than to the STN input stimulation (**Figure 5e**). Similarly, fewer PV^+^ neurons (73.3%, 11 out of 15 neurons) and Npas1^+^ neurons (54.4%, 6 out of 11 neurons) responded to PPN input stimulation (PV^+^ = 54.75 ± 30.5 pA, n = 11 neurons; Npas1^+^ = 54.32 ± 33.2 pA, n = 6 neurons) than to STN input stimulation (**Figure 5e**). Unlike the strong neuron subtype-bias in the strength of input from the STN, there was no statistical difference in the strength of inputs from PF Thal or PPN to the two GPe neuron subpopulations (PF Thal: n_PV_ = 19 neurons, n_Npas1_ = 15 neurons, P = 0.54; PPN: n_PV_ = 11 neurons, n_Npas1_ = 6 neurons, P = 0.35) (**Figure 5e**).

Pharmacological dissection of the synaptic responses in both PV^+^ neurons and Npas1^+^ neurons confirmed that the transmission at STN-GPe synapses was mediated by ionotropic glutamate receptors (**Figure 6a**) as suggested by earlier studies (Kita and Kitai, 1991; Kita, 1992). Optogenetic stimulation of STN terminals in the presence of an NMDA receptor antagonist CPP (10 µM) resulted in responses with only a fast component (not shown); CPP (10 µM) and NBQX (5 µM) completely eliminated the synaptic responses. These results were consistent with prior ultrastructural studies showing that STN terminals form AMPA and NMDA receptor-containing synapses on dendrites of GPe neurons (Bernard and Bolam, 1998; Clarke and Bolam, 1998; Koshimizu et al., 2013). Our experiments collectively showed that the STN provides the primary excitatory drive to GPe neurons, and that it is unique in its cell-targeting properties.

**Figure 6.**
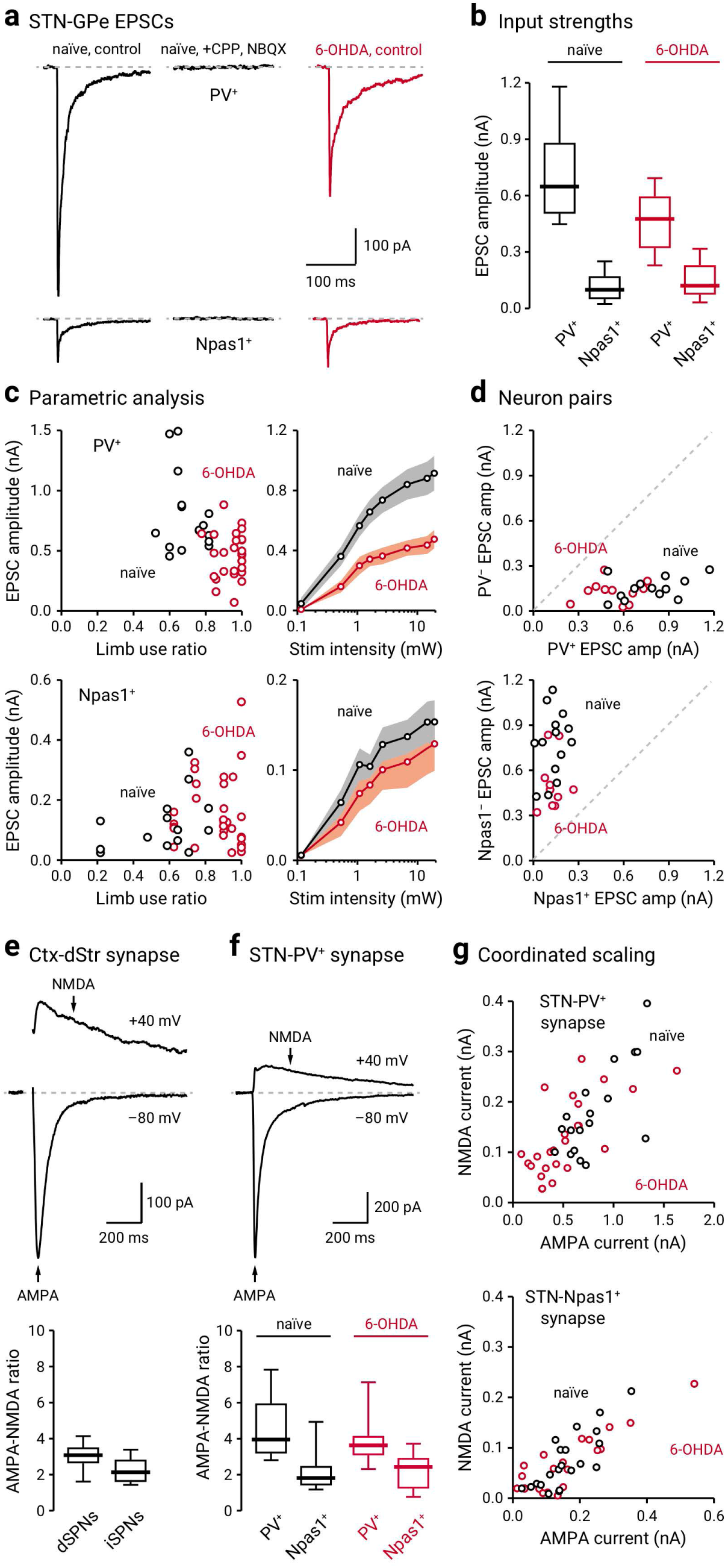
STN-GPe input is weakened following chronic 6-OHDA lesion. **a.** Left: Representative voltage-clamped recordings of a PV^+^ neuron (top) and a Npas1^+^ neuron (bottom) in naïve mice (black) showing that optogenetic stimulation of STN terminals evoked EPSCs. Middle: Application of CPP (10 µM) and NBQX (5 µM) completely eliminated the evoked EPSCs. Right: Representative EPSC recordings from a voltage-clamped PV^+^ neuron and a Npas1^+^ neuron in a chronic 6-OHDA lesioned mouse (red). **b.** Population data showing EPSC amplitudes measured from PV^+^ neurons and Npas1^+^ neurons. Data from naïve (black) and chronic 6-OHDA lesioned (red) mice are shown. **c.** Left: EPSC amplitudes of PV^+^ neurons (top) or Npas1^+^ neurons (bottom) were plotted against limb use ratio, which provides a measure of the extent of the lesion. Each marker indicates a cell. Right: Input-output curves from EPSCs measured from PV^+^ neurons (top) and Npas1^+^ neurons (bottom). Each circle represents the mean EPSC amplitude measured at a particular light intensity, and the shaded area indicates standard error of the mean. **d.** Top: EPSC amplitudes measured from neighboring (within 150 µm apart) PV^+^ neurons and PV^−^ neurons in naïve (black) and chronic 6-OHDA lesioned (red) PV-L-tdTom mice. Bottom: EPSC amplitudes measured from neighboring Npas1^+^ neurons and Npas1^−^ neurons in naïve (black) and chronic 6-OHDA lesioned (red) Npas1-Cre-tdTom mice. Each marker represents a pair of positively- and negatively-identified neurons recorded from the same field. The dashed line represents the unity line. **e.** Top: Representative synaptic responses from a voltage-clamped spiny projection neuron (SPN) in the dStr. Corticostriatal (Ctx-Str) EPSCs were evoked with electrical stimulation; stimulus artifacts are not displayed. The gray line represents the baseline. Neurons were voltage-clamped at –80 mV and +40 mV to measure AMPA and NMDA receptor-dependent currents, respectively. The stimulation artifact was removed for clarity. Bottom: Population data for AMPA-NMDA ratio in direct-pathway spiny projection neurons (dSPNs) and indirect-pathway spiny projection neurons (iSPNs). **f.** Representative synaptic responses from a voltage-clamped PV^+^ neuron. EPSCs were measured with optogenetic stimulation of STN input. Note the relatively small NMDA current in the STN-PV^+^ input. Bottom: Population data for AMPA-NMDA ratio in PV^+^ neurons and Npas1^+^ neurons with stimulation of STN terminals in naïve (black) and chronic 6-OHDA lesioned (red) mice. **g.** The relationship between NMDA current and AMPA current in PV^+^ neurons (top) and Npas1^+^ neurons (bottom) (with stimulation of STN input) in naïve (black) and chronic 6-OHDA lesioned (red) mice. Each marker represents a cell.

It is established that PV^+^ neurons cluster in the dorsolateral regions of the GPe (Hernandez et al., 2015; Saunders et al., 2016; Karube et al., 2019; Abecassis et al., 2020). In addition, anatomical tracing studies indicate that the STN input to the GPe is topographically organized (Iwamuro, 2011; Nambu, 2011). It is possible that the observed differences in the measured strength of input from STN to PV^+^ neurons and to Npas1^+^ neurons were due to sampling bias across different spatial subdomains within the GPe. To this end, neighboring tdTomato^+^ and tdTomato^−^ neurons (less than 150 µm apart) in both PV-L-tdTom (PV-Cre;LSL-tdTom) and Npas1-Cre-tdTom mice were sampled. EPSC amplitudes in PV^+^ neurons were consistently larger than those in PV^−^ neurons (PV^+^: 674.01 ± 174.47 pA, n = 34 neurons; PV^−^: 136.38 ± 60.94 pA, n = 12 neurons, P < 0.0001) (**Figure 6d**). On the other hand, Npas1^+^ neurons had smaller EPSC amplitudes than Npas1^−^ neurons (Npas1^+^: 128.30 ± 63.05 pA, n = 41 neurons; Npas1^−^: 784.82 ± 191.25 pA, n = 16 neurons; P < 0.0001) (**Figure 6d**).

### STN-PV^+^ input is weakened in a chronic model of PD

The STN-GPe network shows abnormally synchronized oscillations in both patients and animal models of PD (Hurtado et al., 1999; Plenz and Kital, 1999; Brown, 2006). Critically, both STN lesioning and deep brain stimulation abolish these pathological oscillatory activities and have profound therapeutic benefits in alleviating the motor symptoms of PD (Bergman et al., 1990; Hurtado et al., 1999; Vitek et al., 2004; Brown, 2006; Kuhn et al., 2006; Vitek et al., 2012). Despite the clinical importance of the STN-GPe network, biophysical description of the alterations of the STN-GPe input in PD remains to be established. To this end, we examined the STN input to PV^+^ neurons and Npas1^+^ neurons in the chronic 6-OHDA lesion model of PD. Similar to the observations in naïve mice, the STN input to PV^+^ neurons was stronger than that to Npas1^+^ neurons in chronic 6-OHDA lesioned mice, as measured by EPSC amplitude. Importantly, STN input to PV^+^ neurons was selectively reduced in chronic 6-OHDA lesioned mice (469.12 ± 154.33 pA, n = 36 neurons, P < 0.0001) (**Figure 6a–d**); this difference was observed across a range of stimulation intensities. On the contrary, STN input to Npas1^+^ neurons did not show a detectable difference (113.28 ± 58.47 pA, n = 37 neurons, P = 0.91) (**Figure 6a–d**).

The neuronal makeup of the STN is generally thought to be homogeneous (Yelnik and Percheron, 1979; Hammond and Yelnik, 1983; Saunders et al., 2016). However, recent studies show that STN neurons are more heterogeneous than previously expected (Sato et al., 2000b; Koshimizu et al., 2013; Xiao et al., 2015; Papathanou et al., 2019). Although single axons can display target cell-specific properties (Shigemoto et al., 1996; Patel et al., 2013; Sun et al., 2018), an alternative explanation for the differences in the STN input to PV^+^ neurons and Npas1^+^ neurons in naïve mice and their alterations in chronic 6-OHDA lesioned mice could be due to cell-specific alterations in postsynaptic receptor properties. Therefore, we biophysically-isolated AMPA and NMDA receptor-dependent currents to measure their relative contribution to the synaptic responses to STN input in PV^+^ neurons and Npas1^+^ neurons. In addition, we compared these AMPA and NMDA measurements to those of a well-studied glutamatergic synapse—the corticostriatal synapse (**Figure 6e & f**). In naïve mice, both AMPA and NMDA receptor-mediated EPSCs in PV^+^ neurons were larger than those in Npas1^+^ neurons (AMPA current: PV^+^ = 685.06 ± 155.03 pA, n = 22 neurons; Npas1^+^ = 141.11 ± 45.53 pA, n = 18 neurons; P < 0.0001; NMDA current: PV^+^ = 172.30 ± 54.26 pA, n = 22 neurons; Npas1^+^ = 62.19 ± 38.39 pA, n = 18 neurons, P < 0.0001) (**Figure 6f & g**). The AMPA-NMDA ratio of PV^+^ neurons was also larger than that in Npas1^+^ neurons (PV^+^ = 3.96 ± 0.82, n = 22 neurons; Npas1^+^ = 1.82 ± 0.45, n = 18 neurons; P < 0.0001). The AMPA-NMDA ratio of the STN-PV^+^ input was larger than that observed at the corticostriatal synapse (dSPN = 3.09 ± 0.40, n = 10 neurons, P = 0.014; iSPN = 2.14 ± 0.52, n = 16 neurons; P < 0.0001) (**Figure 6e & f**). The difference in the AMPA-NMDA ratio in PV^+^ neurons and Npas1^+^ neurons indicates that different receptor complements mediate the transmission.

Both AMPA and NMDA receptor-mediated currents in PV^+^ neurons were reduced in chronic 6-OHDA lesioned mice compared to naïve mice (AMPA current = 430.79 ± 150.51 pA, n = 27 neurons, P = 0.00030; NMDA current = 116.94 ± 54.69 pA, n = 27 neurons, P = 0.024) (**Figure 6f & g**). This appeared to be a coordinated regulation, as the AMPA-NMDA ratio was unchanged in chronic 6-OHDA lesioned mice (PV^+^ = 3.62 ± 0.55, n = 27 neurons, P = 0.18) (**Figure 6f & g**). In contrast with the findings in PV^+^ neurons, AMPA and NMDA receptor-mediated currents were unchanged in Npas1^+^ neurons following chronic 6-OHDA lesion (AMPA current = 122.07 ± 52.31 pA, n = 17 neurons, P = 0.73; NMDA current = 84.35 ± 33.69 pA, n = 17 neurons, P = 0.81) (**Figure 6f & g**).

Given the decrease in the functional connectivity of the STN-PV^+^ input, we expect to see a correlated anatomical alteration following chronic 6-OHDA lesion. Contrary to our prediction, we did not find an expected decrease in the density of STN axonal fibers in the GPe following the chronic 6-OHDA lesion (naïve, lateral = 3.61 ± 0.23 x10^7^ a.u., n = 6 mice; 6-OHDA, lateral = 3.89 ± 0.63 x10^7^ a.u., n = 7 mice; P = 0.23; naïve, intermediate = 3.80 ± 0.90 x10^7^ a.u., n = 6 mice; 6-OHDA, intermediate = 4.21 ± 0.26 x10^7^ a.u., n = 8 mice; P = 0.75; naïve, medial = 2.40 ± 0.88 x10^7^ a.u., n = 6 mice; 6-OHDA, medial = 2.66 ± 0.47 x10^7^ a.u., n = 8 mice; P = 0.66) (**Figure 7a & b**). As the density of the STN-GPe axons may not be tightly associated with the number of release sites, we then examined the abundance of putative STN-GPe boutons. By measuring density of eYFP-labeled puncta that were immunopositive for VGluT2, we found a decrease in the density of STN presynaptic boutons in the GPe following chronic 6-OHDA lesion (naïve = 5.1 ± 0.98 count/µm^3^, n = 15 sections; 6-OHDA = 3.6 ± 1.2 count/µm^3^, n = 17 sections; P = 0.028) (**Figure 7c**). This finding corroborated the decrease in both AMPA and NMDA receptor currents in PV^+^ neurons (**Figure 6g**). Although alterations in the release probability and quantal properties of the STN-GPe synapse cannot be excluded, our data collectively show a loss of STN-GPe connectivity in a chronic model of PD.

**Figure 7.**
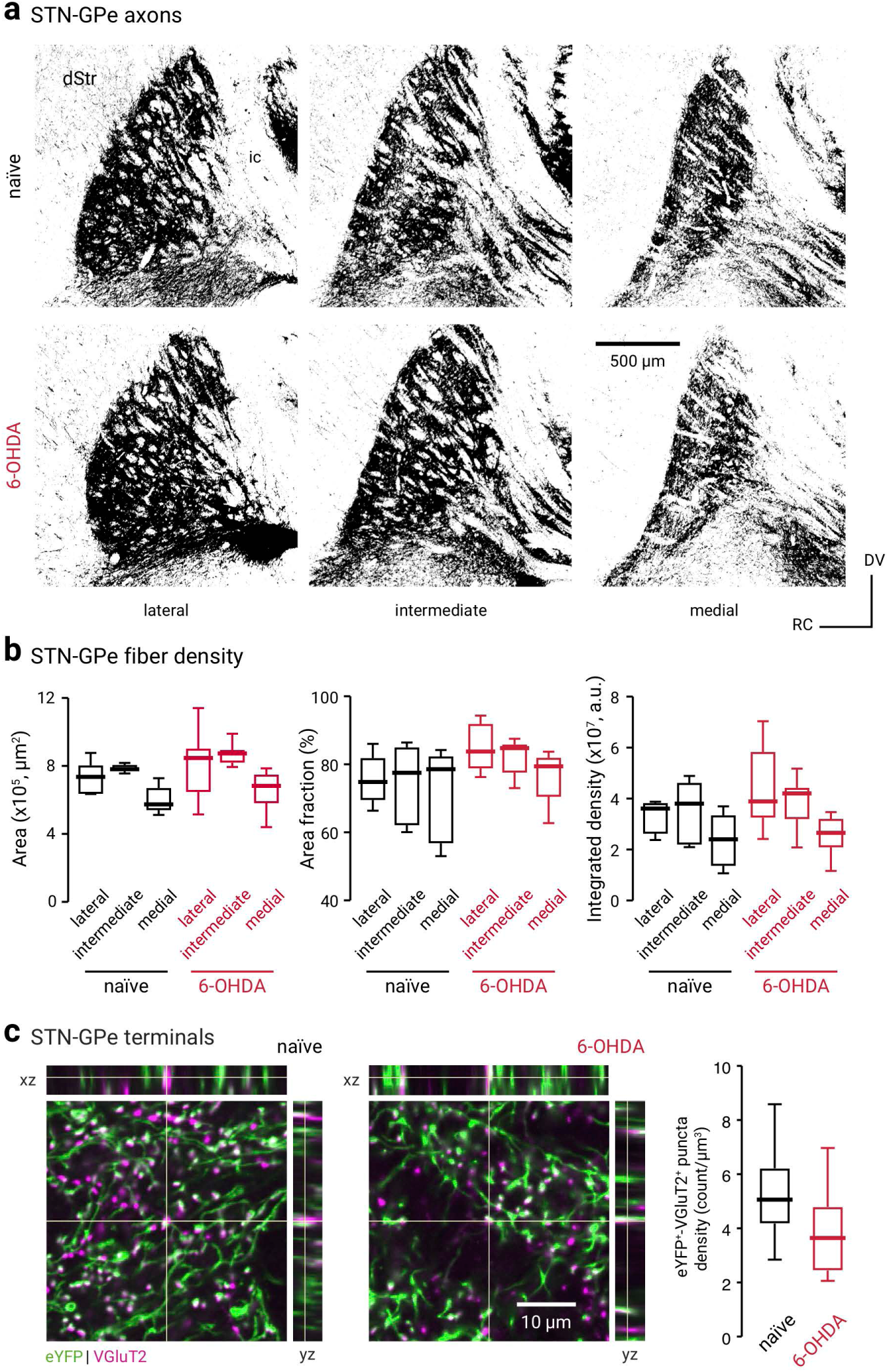
The cell-type specificity of STN-GPe EPSCs is not due to topographical biasing. **a.** Confocal micrographs showing the GPe in sagittal brain sections of a naïve (top, black) or chronic 6-OHDA lesioned (bottom, red) C57BL/6J mice with ChR2-eYFP-expressing AAVs in the STN. Lateral, intermediate, and medial sections of the GPe are shown. Abbreviations: dStr, dorsal striatum; ic, internal capsule. **b.** Analysis of the eYFP signal in naïve (black) and chronic 6-OHDA lesioned (red) mice. The surface area of the GPe (left), percent area covered by the eYFP signal (middle), and the integrated density of the eYFP signal (right) are quantified. **c.** Left & middle: Confocal micrographs showing the density of eYFP- and VGluT2-immunoreactive elements in the GPe from naïve and chronic 6-OHDA lesioned mice. Breakout panels show orthogonal xz-projection (top) and yz-projection (right). Crosshairs indicate the pixel of interest. The colocalization of the signals is shown as white. Right: Box plots summarize the density of putative STN-GPe terminals in naïve (black) and chronic 6-OHDA lesioned (red) mice.

In light of the difference in the STN input to the two GPe neuron classes, we examined whether STN input also causes distinct changes in the activity of PV^+^ neurons and Npas1^+^ neurons. We monitored the firing of GPe neurons in response to optogenetic stimulation of the STN input. Consistent with our prior work (Hernandez et al., 2015; Abecassis et al., 2020), PV^+^ neurons and Npas1^+^ neurons have distinct basal activity levels (PV^+^: baseline_naïve_ = 18.26 ± 3.74 Hz, n = 23 neurons; Npas1^+^: baseline_naïve_ = 8.91 ± 2.97 Hz, n = 10 neurons; P < 0.0001) (**Figure 8a & b**). In response to optogenetic stimulation of STN input, both PV^+^ neurons and Npas1^+^ neurons showed increases in their firing (PV^+^: stim_naïve_ = 51.48 ± 6.93 Hz, n = 23 neurons, P < 0.0001; Npas1^+^: stim_naïve_ = 31.68 ± 4.95 Hz, n = 10 neurons, P = 0.0020) (**Figure 8a & b**). In naïve mice, the fold change in the firing of PV^+^ neurons and Npas1^+^ neurons with STN stimulation was not different (PV^+^_naïve_: +3.09 ± 0.90 fold, n = 23 neurons; Npas1^+^_naïve_: +3.87 ± 1.13 fold, n = 10 neurons; P = 0.48). In 6-OHDA lesion, optogenetic stimulation of STN input also resulted in increases in the firing of PV^+^ neurons and Npas1^+^ neurons (PV^+^: baseline_6-OHDA_ = 20.79 ± 2.97 Hz, stim_6-OHDA_ = 50.49 ± 9.90 Hz, n = 15 neurons, P < 0.0001; Npas1^+^: baseline_6-OHDA_ = 5.94 ± 1.98 Hz, stim_6-OHDA_ = 16.83 ± 4.95 Hz, n = 11 neurons, P = 0.0010). Consistent with a weakening of the STN-PV^+^ input, PV^+^ neurons showed a selective reduction in the fold-change of firing following a chronic 6-OHDA lesion (PV^+^: stim_naïve_ = +3.09 ± 0.90 fold, n = 23 neurons; stim_6-OHDA_ = +2.38 ± 0.27 fold, n = 15 neurons; P = 0.0049) (**Figure 8a & b**). In contrast, Npas1^+^ neurons did not show a change in the fold-change in their firing between naïve and chronic 6-OHDA lesion (Npas1^+^: stim_naïve_ = +3.87 ± 1.13 fold, n = 10 neurons; stim_6-OHDA_ = +2.98 ± 0.96 fold, n = 11 neurons; P = 0.24).

**Figure 8.**
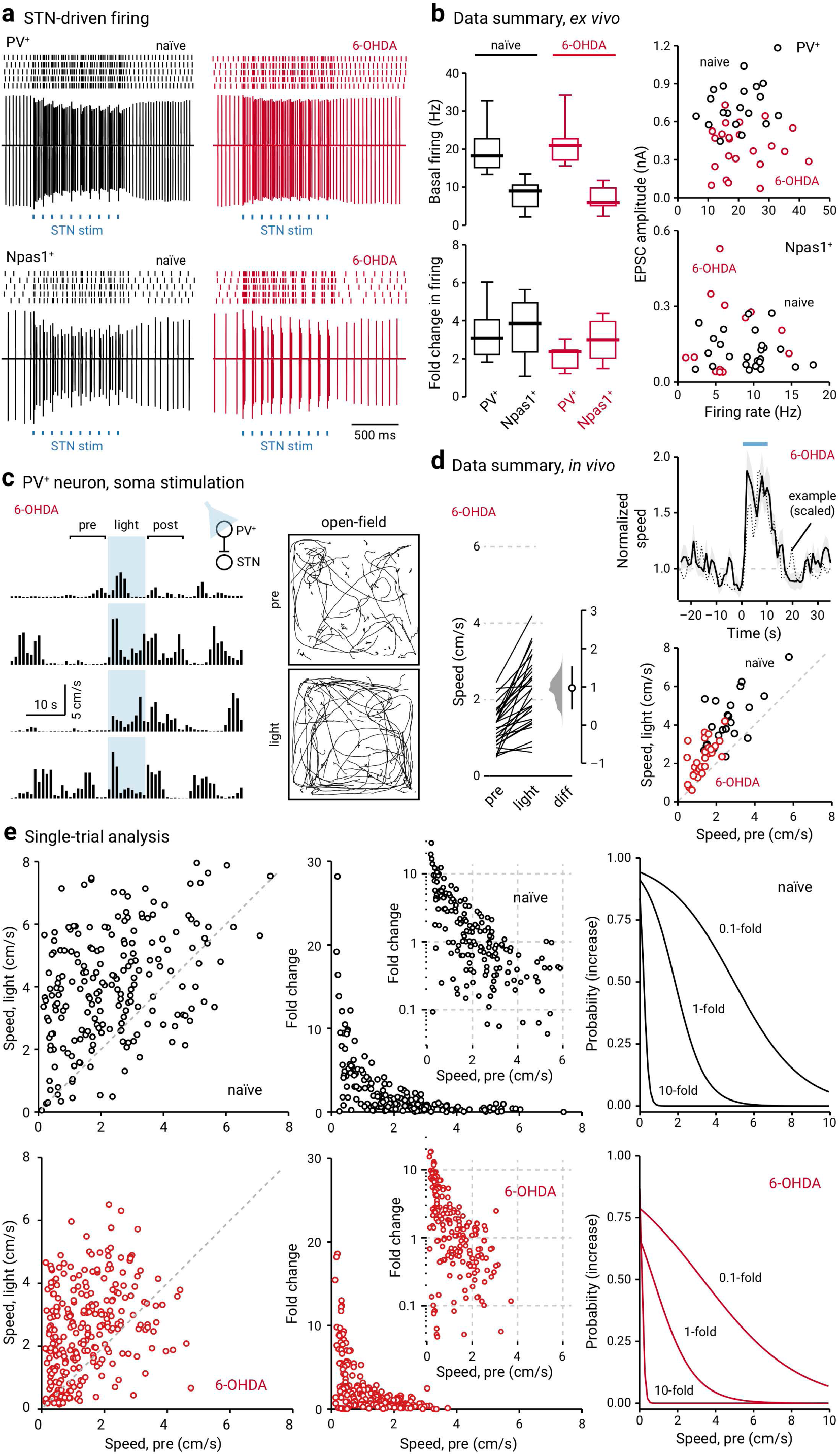
PV^+^ neuron stimulation lessens hypokinetic symptoms. **a.** Top: Representative cell-attached recordings from PV^+^ neurons in naïve (left, black) and chronic 6-OHDA lesioned (right, red) mice. Raster plots show trials from a single PV^+^ neuron, where each raster corresponds to an action potential. Bottom: Representative cell-attached recordings from Npas1^+^ neurons in naïve (left, black) and 6-OHDA lesion (right, red) mice. Each blue square represents a 2-ms light pulse delivered to the GPe. **b.** Left: Box plots summarizing the population data from naïve (left, black) and chronic 6-OHDA lesioned (right, red) mice. Right: Scatter plots showing the relationship between EPSC amplitude and spontaneous firing rate of PV^+^ neurons (top) and Npas1^+^ neurons (bottom). Data from both naïve (black) and chronic 6-OHDA lesioned (red) mice are displayed. Each marker indicates a cell. **c.** Open field activity in response to optogenetic stimulation of PV^+^ neurons in the GPe in chronic 6-OHDA lesioned mice. Left: A representative example of locomotion activity of a chronic 6-OHDA lesioned mouse across four individual trials. The schematic diagram shows the cell type and site of light delivery. ‘Pre-period’ indicates 10 s before light delivery; ‘post-period’ indicates 10 s after light delivery. Blue shaded area indicates the duration (10 s) of light delivery. Right: Movement tracks corresponding to the ‘pre-period’ (top) and ‘light-period’ (bottom). Six representative mice (ten trials from each) are presented. **d.** Left: Slopegraph showing the response of each mouse upon optogenetic stimulation of PV^+^ neurons in chronic 6-OHDA lesioned mice. The slope of each line shows the magnitude and direction of change. Median difference is plotted on a floating axis. The smoothed density represents bootstrap resampled data. The median difference is indicated as a circle and the 95% confidence interval is indicated by the length of vertical lines. Top, right: A plot showing the relationship between normalized speed and time. Blue bar indicates the duration (10s) of light delivery. The dotted horizontal line indicates the normalized baseline motor activity level. Black solid trace is the average distance from all mice; the shaded area shows the standard error of the mean. Black dotted trace is a representative example from a single mouse; data were scaled to facilitate comparison. Bottom, right: Speed during ‘light-period’ against speed during ‘pre-period’. Data from chronic 6-OHDA lesioned PV-Cre mice expressing ChR2-eYFP are displayed. Each marker represents a mouse. The diagonal line indicates unity (i.e., x = y). 6-OHDA data points are systematically shifted upwards relative to unity. Locomotor activity was higher with light delivery. Naïve data (black) from Figure 1b are replotted for comparison. **e.** Left: Scatter plots showing the pairwise relationship between speed during ‘light-period’ and speed during ‘pre-period’ in naïve (top) and chronic 6-OHDA lesioned PV-Cre mice are shown (bottom). To facilitate comparison, data from naïve mice are replotted (top). The diagonal line indicates unity. Middle: Fold change in speed with light delivery against speed during ‘pre-period’ from PV-Cre mice expressing ChR2-eYFP. Each marker represents a trial. Inset: Same data displayed with fold increase on y-axis on a log scale. Right: Logistic regression curves fitted to data for 0.1-fold, 1-fold, and 10-fold increases are displayed.

### Stimulation of PV^+^ neurons lessens hypokinetic symptoms

In agreement with the established relationship between STN activity and movement suppression (Hamani et al., 2004; Aron and Poldrack, 2006; Aron et al., 2007; Eagle et al., 2008; Schmidt et al., 2013; Schweizer et al., 2014; Fife et al., 2017; Wessel and Aron, 2017; Adam et al., 2020), both direct recordings and theoretical models assert that the hypokinetic symptoms of PD are a result of excessive STN activity (Albin et al., 1989; Bergman et al., 1994; Wichmann and DeLong, 1996; Bergman et al., 1998; Obeso et al., 2000; DeLong and Wichmann, 2007; Zaidel et al., 2009; Sharott et al., 2014; DeLong and Wichmann, 2015; McGregor and Nelson, 2019). Importantly, lesioning and inactivation studies from animal models of PD further support this idea (Levy et al., 2001; Yoon et al., 2014), but see (McIver et al., 2019). In other words, the weakening of the STN-PV^+^ input that we observed in the chronic 6-OHDA lesioned mice *ex vivo* can thus be a form of homeostatic scaling in response to the increased excitatory drive. However, this alteration may, in fact, be maladaptive as it would lead to decreased inhibitory output from the GPe to the STN. The resultant increased STN activity would in turn lead to motor suppression. If our interpretation is correct, then optogenetic stimulation of PV^+^ neurons, which would inhibit STN neurons, should restore motor activity in chronic 6-OHDA lesioned mice. Chronic 6-OHDA lesioned mice have a reduced basal ambulatory activity (naïve = 2.62 ± 0.86 cm/s, n = 12 mice; 6-OHDA = 1.72 ± 0.38 cm/s, n = 14 mice, P = 0.011), as expected. Under this condition, ChR2-mediated stimulation of PV^+^ neurons increased locomotion in chronic 6-OHDA mice (patterned: +0.48 ± 0.26 fold, n = 14 mice, P = 0.00037; sustained: +0.72 ± 0.20 fold, n = 14 mice, P = 0.00012); the extent of the increase was comparable to that observed in naïve mice (patterned: P = 0.89; sustained: P = 0.82) (**Figure 8c–d**). Moreover, stimulation of PV^+^ neurons in 6-OHDA mice induced motor responses similar to the spontaneous ambulatory activity in naïve mice (naïve_pre_ = 2.62 ± 0.86 cm/s, n = 12 mice; 6-OHDA_light_ = 2.57 ± 0.69 cm/s, n = 14 mice, P = 0.22). As shown in **Figure 8e**, although the net locomotor activity during the light-period was distinct from its pre-period level in chronic 6-OHDA lesioned mice, there was an increased number of trials that showed a negative change (i.e., data points that are below unity in the x-y scatter plots) in locomotion following the chronic 6-OHDA lesion (naïve: 44 out of 230 trials, 19.1%; 6-OHDA: 76 out of 280 trials, 27.1%; P = 0.036). Furthermore, the strong negative relationship between pre-period speed and the increase in locomotion that was observed in naïve mice was more pronounced in chronic 6-OHDA lesioned mice. As described by a logistic regression function, the probability to observe an increased locomotion is reduced following the chronic 6-OHDA lesion (**Figure 8e**, **Table 4**).

It has been shown previously that the stimulation of PV^+^ neurons produced persistent motor promotion in an acute 6-OHDA lesion model (Mastro et al., 2017). However, it was unknown whether the lasting response can be generalized to the chronic 6-OHDA lesion model. Here, we examined whether the motor effects induced by optogenetic stimulation in PV^+^ neurons were a function of previous stimulation history. As illustrated in **Figure 9a**, while motor promotion was readily observed (in both naïve and chronic 6-OHDA lesioned mice) when PV^+^ neurons were optogenetically stimulated, the magnitude of the effect was not associated with the order of the trials. This was determined by examining the trial number as a predictor variable in a logistic regression model. Critically, this analysis shows that repetitive optogenetic stimulation of PV^+^ neurons did not lead to a sustained elevation of locomotor activity in the chronic 6-OHDA lesioned mice. This inference was confirmed by extending the time window for monitoring motor activity before and after the entire set of optogenetic stimuli in a subset of mice; the locomotion speed was maintained at the same level before and after the stimuli (pre_5min_: 0.96 ± 0.29 cm/s, post_5min_ : 1.07 ± 0.15 cm/s, n = 4, P = 0.89) (**Figure 9b**). Lastly, a summary of motor effects under different conditions is tabulated in **Table 5**.

**Figure 9.**
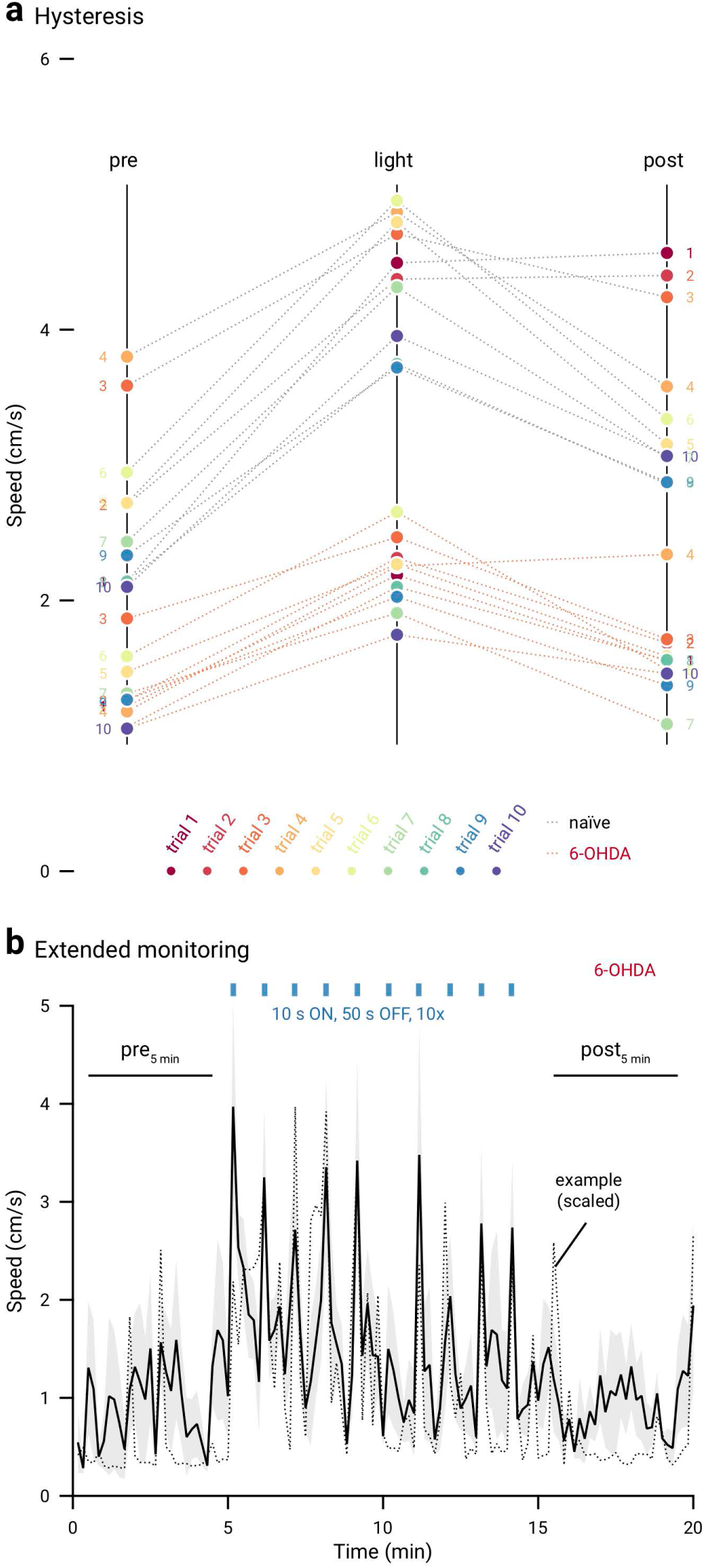
No persistent motor effects are induced by *in vivo* optogenetic stimulation in chronic 6-OHDA mice. **a.** Slopegraph showing the size and reversibility of motor responses induced by optogenetic stimulation of PV^+^ neurons in naïve (gray dotted lines) and chronic 6-OHDA lesioned mice (red dotted lines). Ten trials (circles) were run on each mouse. Trial numbers are denoted by colors (rainbow). Data from sustained (naïve: n = 12; 6-OHDA: n = 14) and patterned (naïve: n = 12; 6-OHDA: n = 14) stimulation are combined. The slope of each line shows the magnitude and direction of change. **b.** A plot showing the motor activity of a subset of chronic 6-OHDA lesioned mice (n = 4) across time. Blue bar indicates the duration of light delivery; ten stimuli were delivered. Black solid trace is the average distance from all mice; the shaded area shows the standard error of the mean. Black dotted trace is a representative example from a single mouse; data were scaled to facilitate comparison.

**Table 5.**
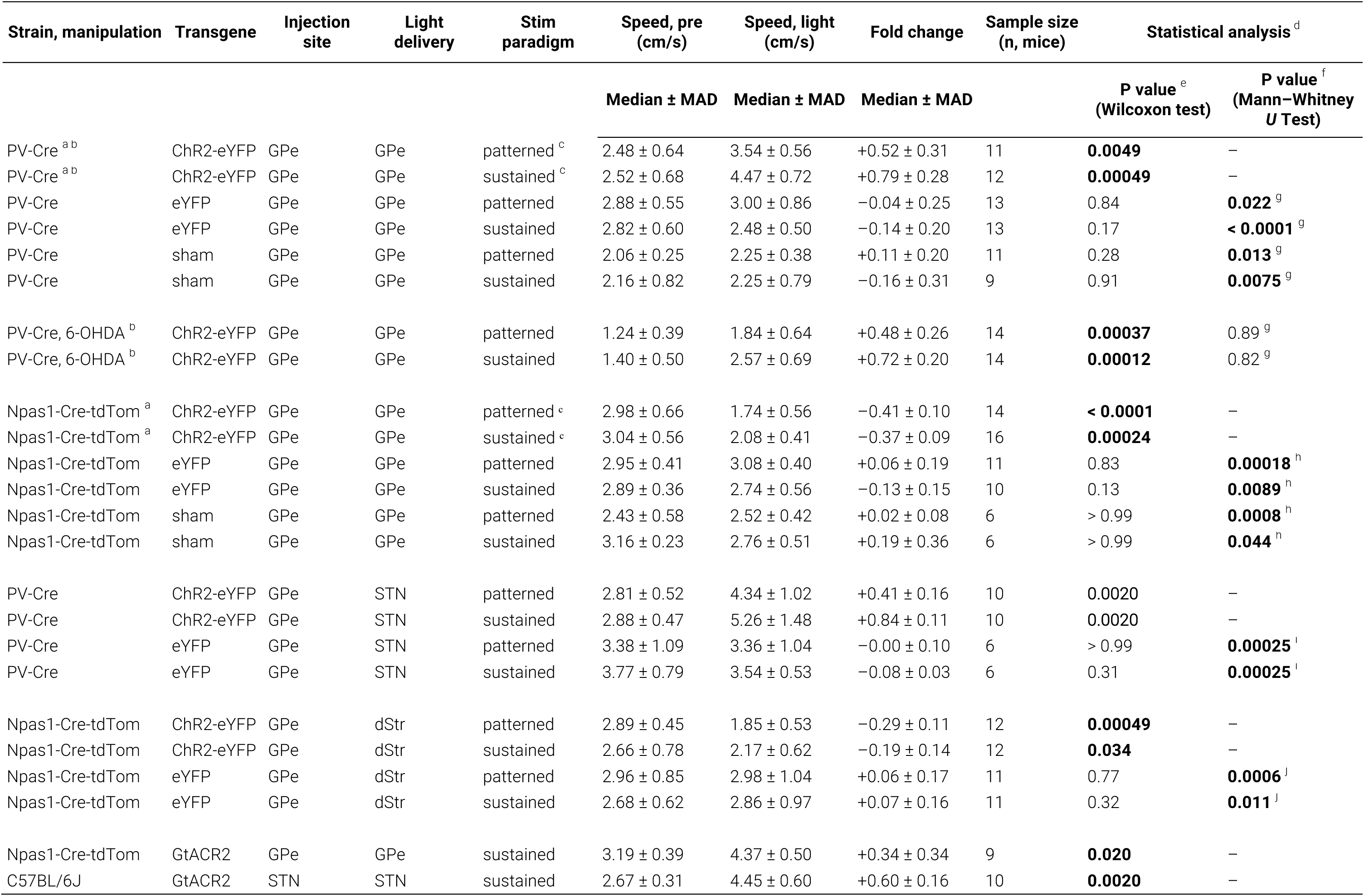
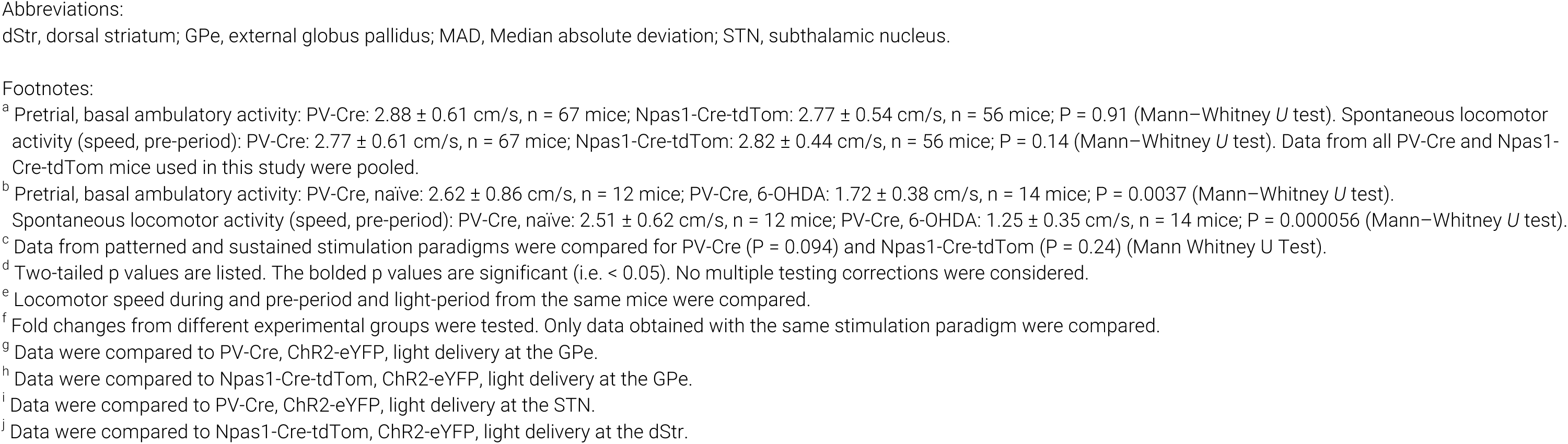
Statistical summary for behavioral analyses.

| Strain, manipulation | Transgene | Injection site | Light delivery | Stim paradigm | Speed, pre (cm/s) | Speed, light (cm/s) | Fold change | Sample size (n, mice) | Statistical analysis<sup>d</sup> |  |
| --- | --- | --- | --- | --- | --- | --- | --- | --- | --- | --- |
|  |  |  |  |  | Median ± MAD | Median ± MAD | Median ± MAD |  | P value<sup>e</sup><br>(Wilcoxon test) | P value<sup>f</sup><br>(Mann–Whitney U Test) |
| PV-Cre<sup>a,b</sup> | ChR2-eYFP | GPe | GPe | patterned<sup>c</sup> | 2.48 ± 0.64 | 3.54 ± 0.56 | +0.52 ± 0.31 | 11 | **0.0049** | – |
| PV-Cre<sup>a,b</sup> | ChR2-eYFP | GPe | GPe | sustained<sup>c</sup> | 2.52 ± 0.68 | 4.47 ± 0.72 | +0.79 ± 0.28 | 12 | **0.00049** | – |
| PV-Cre | eYFP | GPe | GPe | patterned | 2.88 ± 0.55 | 3.00 ± 0.86 | –0.04 ± 0.25 | 13 | 0.84 | **0.022<sup>g</sup>** |
| PV-Cre | eYFP | GPe | GPe | sustained | 2.82 ± 0.60 | 2.48 ± 0.50 | –0.14 ± 0.20 | 13 | 0.17 | **< 0.0001<sup>g</sup>** |
| PV-Cre | sham | GPe | GPe | patterned | 2.06 ± 0.25 | 2.25 ± 0.38 | +0.11 ± 0.20 | 11 | 0.28 | **0.013<sup>g</sup>** |
| PV-Cre | sham | GPe | GPe | sustained | 2.16 ± 0.82 | 2.25 ± 0.79 | –0.16 ± 0.31 | 9 | 0.91 | **0.0075<sup>g</sup>** |
| PV-Cre, 6-OHDA<sup>b</sup> | ChR2-eYFP | GPe | GPe | patterned | 1.24 ± 0.39 | 1.84 ± 0.64 | +0.48 ± 0.26 | 14 | **0.00037** | 0.89<sup>g</sup> |
| PV-Cre, 6-OHDA<sup>b</sup> | ChR2-eYFP | GPe | GPe | sustained | 1.40 ± 0.50 | 2.57 ± 0.69 | +0.72 ± 0.20 | 14 | **0.00012** | 0.82<sup>g</sup> |
| Npas1-Cre-tdTom<sup>a</sup> | ChR2-eYFP | GPe | GPe | patterned<sup>c</sup> | 2.98 ± 0.66 | 1.74 ± 0.56 | –0.41 ± 0.10 | 14 | **< 0.0001** | – |
| Npas1-Cre-tdTom<sup>a</sup> | ChR2-eYFP | GPe | GPe | sustained<sup>c</sup> | 3.04 ± 0.56 | 2.08 ± 0.41 | –0.37 ± 0.09 | 16 | **0.00024** | – |
| Npas1-Cre-tdTom | eYFP | GPe | GPe | patterned | 2.95 ± 0.41 | 3.08 ± 0.40 | +0.06 ± 0.19 | 11 | 0.83 | **0.00018<sup>h</sup>** |
| Npas1-Cre-tdTom | eYFP | GPe | GPe | sustained | 2.89 ± 0.36 | 2.74 ± 0.56 | –0.13 ± 0.15 | 10 | 0.13 | **0.0089<sup>h</sup>** |
| Npas1-Cre-tdTom | sham | GPe | GPe | patterned | 2.43 ± 0.58 | 2.52 ± 0.42 | +0.02 ± 0.08 | 6 | > 0.99 | **0.0008<sup>h</sup>** |
| Npas1-Cre-tdTom | sham | GPe | GPe | sustained | 3.16 ± 0.23 | 2.76 ± 0.51 | +0.19 ± 0.36 | 6 | > 0.99 | **0.044<sup>h</sup>** |
| PV-Cre | ChR2-eYFP | GPe | STN | patterned | 2.81 ± 0.52 | 4.34 ± 1.02 | +0.41 ± 0.16 | 10 | **0.0020** | – |
| PV-Cre | ChR2-eYFP | GPe | STN | sustained | 2.88 ± 0.47 | 5.26 ± 1.48 | +0.84 ± 0.11 | 10 | **0.0020** | – |
| PV-Cre | eYFP | GPe | STN | patterned | 3.38 ± 1.09 | 3.36 ± 1.04 | –0.00 ± 0.10 | 6 | > 0.99 | **0.00025<sup>i</sup>** |
| PV-Cre | eYFP | GPe | STN | sustained | 3.77 ± 0.79 | 3.54 ± 0.53 | –0.08 ± 0.03 | 6 | 0.31 | **0.00025<sup>i</sup>** |
| Npas1-Cre-tdTom | ChR2-eYFP | GPe | dStr | patterned | 2.89 ± 0.45 | 1.85 ± 0.53 | –0.29 ± 0.11 | 12 | **0.00049** | – |
| Npas1-Cre-tdTom | ChR2-eYFP | GPe | dStr | sustained | 2.66 ± 0.78 | 2.17 ± 0.62 | –0.19 ± 0.14 | 12 | **0.034** | – |
| Npas1-Cre-tdTom | eYFP | GPe | dStr | patterned | 2.96 ± 0.85 | 2.98 ± 1.04 | +0.06 ± 0.17 | 11 | 0.77 | **0.0006<sup>j</sup>** |
| Npas1-Cre-tdTom | eYFP | GPe | dStr | sustained | 2.68 ± 0.62 | 2.86 ± 0.97 | +0.07 ± 0.16 | 11 | 0.32 | **0.011<sup>j</sup>** |
| Npas1-Cre-tdTom | GtACR2 | GPe | GPe | sustained | 3.19 ± 0.39 | 4.37 ± 0.50 | +0.34 ± 0.34 | 9 | **0.020** | – |
| C57BL/6J | GtACR2 | STN | STN | sustained | 2.67 ± 0.31 | 4.45 ± 0.60 | +0.60 ± 0.16 | 10 | **0.0020** | – |

Abbreviations:
dStr, dorsal striatum; GPe, external globus pallidus; MAD, Median absolute deviation; STN, subthalamic nucleus.

## Discussion

In this study, by optogenetically manipulating the activity of specific GPe neuron subpopulations and their downstream targets, we concluded that PV^+^ neurons are movement-promoting and Npas1^+^ neurons are movement-inhibiting. Consistent with their distinct functional roles, *ex vivo* electrophysiology further delineated that the STN input biased toward PV^+^ neurons. By using a chronic model of PD, we showed that PV^+^ neurons play an important role in the hypokinetic symptoms of the disease. Accordingly, optogenetic stimulation of PV^+^ neurons restored mobility.

### GPe neuron subtypes have opposing roles in motor control

Historically, it has been challenging to assign specific functional roles to the GPe. Lesioning or inactivation studies were largely inconclusive, as these manipulations did not result in a consistent motor phenotype (Norton, 1976; Ossowska et al., 1983; Schneider and Olazabal, 1984; Hauber et al., 1998; Joel et al., 1998; Konitsiotis et al., 1998; Soares et al., 2004; Hegeman et al., 2016). A large number of studies showed that GPe neurons change their activity in relation to movement; however, the identity of the recorded neurons was unknown (DeLong, 1971; Anderson, 1978; Anderson and Horak, 1985; DeLong et al., 1985; Mink and Thach, 1987; Mitchell et al., 1987; Filion et al., 1988; Nambu et al., 1990; Mink and Thach, 1991b, a; Mushiake and Strick, 1995; Parent and Hazrati, 1995; Kimura et al., 1996; Arkadir et al., 2004; Shin and Sommer, 2010; Schmidt et al., 2013; Yoshida and Tanaka, 2016; Gu et al., 2020; Mullie et al., 2020). It has been shown that identified GPe neuron subtypes can display diverse changes in their activity during spontaneous body movements (Dodson et al., 2015). Collectively, these observational studies strongly argue that GPe neurons are involved in motor control. Our data, along with our previous study, unambiguously demonstrate the causality between the activity of GPe neurons and locomotion (Glajch et al., 2016)—and importantly the opposing roles played by PV^+^ neurons and Npas1^+^ neurons. Though we have quantified immobility in this study, we do not have quantitative information about whether optogenetic manipulations may result in altered innate behaviors (such as rearing, grooming, and freezing) that may confound our measurements of locomotion.

Our current data showed that in naïve mice, stimulation of PV^+^ neurons and their terminals in the STN promoted movement. To our knowledge, this has not been previously reported. Consistent with the fact that STN neuron activity is associated with movement inhibition (Hamani et al., 2004; Aron and Poldrack, 2006; Aron et al., 2007; Eagle et al., 2008; Schmidt et al., 2013; Schweizer et al., 2014; Fife et al., 2017; Wessel and Aron, 2017; Adam et al., 2020), we showed that phasic inhibition (either synaptically- or opsin-mediated) of STN neurons is movement-promoting. In addition, as the majority of PV^+^ neurons send bifurcating axons to both the STN and SNr (Bevan et al., 1998; Sato et al., 2000a; Kita, 2007; Fujiyama et al., 2016; Oh et al., 2017); from a circuit organization standpoint, the PV^+^-SNr projection is expected to reinforce the motor effects imposed by the PV^+^-STN projection. Although our observation is in line with the findings from our lab and others (Glajch et al., 2016; Mallet et al., 2016), we do not fully understand the motor suppression produced by Npas1^+^ neurons. While Npas1^+^ neurons have extensive axonal arborization in the dStr, their downstream impact on the excitability of striatal projection neurons is relatively weak (Glajch et al., 2016). It is possible that additional signaling partners are involved.

In this study, we drove the activity of PV^+^ neurons or Npas1^+^ neurons by activating ChR2. These experiments were critical to establishing the causal role of PV^+^ neurons and Npas1^+^ neurons in motor control. By directing light stimulus to different brain structures, we dissected circuit elements that are involved in these motor effects. While these gain-of-function experiments caused observable motor effects, they are insufficient to conclude if the motor effects are the native functions mediated by PV^+^ neurons and Npas1^+^ neurons. We addressed this by performing loss-of-function experiments using GtACR2 to inhibit GPe neuron subtypes or their postsynaptic targets. GPe neurons are known to form intranuclear collaterals (Kita, 1994; Nambu and Llinas, 1997; Mallet et al., 2012; Saunders et al., 2015). As we do not yet have the tools for the selective manipulation of these local collaterals, we will need to rely on *ex vivo* studies to assess their relevance based on the connectivity principle of these local connections. By examining moment-to-moment fluctuations in movement speed, we showed that optogenetically-induced motor responses did not differ from the normal range of spontaneous locomotion displayed by mice. In sum, our *in vivo* studies reinforce the notion from prior *in vivo* electrophysiological studies that GPe neurons are critical for movement.

There are several possible biological benefits to a dual-system of PV^+^ neurons and Npas1^+^ neurons in the GPe. It is conceivable that the two neuron populations with opposing functions form a rheostat in controlling net motor output. However, their functional relationship may not be simply antagonistic. As both initiation and termination of motor programs are equally important and are core functions of the basal ganglia (Redgrave et al., 2010; Jahanshahi et al., 2015), it is possible that PV^+^ neurons and Npas1^+^ neurons are differentially involved in selecting voluntary versus suppressing competing motor programs. Alternatively, GPe neuron subtypes can be selectively engaged in the regulation of antagonistic muscle groups that are necessary for the execution of one specific movement. It is reasonable to think that PV^+^ neurons and Npas1^+^ neurons are both involved in the selection of competing behaviors and the combination of movements that occur either jointly or in succession. Related ideas have been proposed for striatal neurons (Mink, 1996; Klaus et al., 2017; Arber and Costa, 2018; Klaus et al., 2019). To gain further insights into this topic, it will be important in the future to determine the temporal organization of the activity of GPe neuron subtypes in relation to movement.

### The STN forms a reciprocal loop with PV^+^ neurons

In this study, we found that the STN provides a stronger input to PV^+^ neurons than to Npas1^+^ neurons. This finding is supported by a recent *in vivo* electrophysiological study showing STN strongly targets “prototypic” neurons that are predominantly PV^+^ neurons (Ketzef and Silberberg, 2020). This is in contrast with our previous rabies-tracing study that showed a uniform STN input across GPe neuron subtypes (Hunt et al., 2018). In addition, our finding contradicts computational studies that predicted a stronger connection from the STN to a subset of Npas1^+^ neurons (Bogacz et al., 2016; Suryanarayana et al., 2019). Given the known caveats associated with rabies-based tracing (Svoboda, 2019), this discrepancy perhaps is not completely surprising. As we found a notable input from the STN to Npas1^+^ neurons, it is possible that the STN input regulates movement via sending an efference copy to the Npas1^+^ neurons in addition to the primary projection to its downstream target, i.e., PV^+^ neurons. Moreover, the difference in the AMPA-NMDA ratio of the synaptic responses in PV^+^ neurons and Npas1^+^ neurons indicates that the STN input to these neurons is mediated by different complements in the postsynaptic glutamate receptors.

Recent studies indicate the presence of heterogeneity in STN neurons (Sato et al., 2000b; Isoda and Hikosaka, 2008; Koshimizu et al., 2013; Xiao et al., 2015; Papathanou et al., 2019). It is possible that unique STN neurons provide inputs to distinct GPe neuron subpopulations. As we are only beginning to grasp the cellular heterogeneity within both the GPe and STN, additional work is needed to further our understanding of the organization of the STN-GPe network. As PV^+^ neurons form the principal inhibitory innervation to the STN (Mastro et al., 2014; Abdi et al., 2015; Hernandez et al., 2015; Saunders et al., 2016), these data collectively demonstrate a closed reciprocal feedback loop formed between the STN and PV^+^ neurons. Although earlier studies have examined the electrophysiological and anatomical properties of STN-GPe network (Kita and Kitai, 1987, 1991; Nambu et al., 2000; Kita et al., 2004; Kita, 2007; Mastro et al., 2014; Abdi et al., 2015; Hernandez et al., 2015; Saunders et al., 2016; Kovaleski et al., 2020), the cell-type specificity of the STN input in the GPe was not known. Importantly, our new data add critical insights into the cellular constituents that are involved in this reciprocal loop.

In this study, we found that the STN input to PV^+^ neurons is reduced in chronic 6-OHDA lesioned mice. These findings are consistent with the downregulation of glutamate receptors in the GPe of PD models (Porter et al., 1994; Betarbet et al., 2000; Kaneda et al., 2005). We have previously found a decrease in the ambient glutamate content in the GPe following a chronic loss of dopamine (Cui et al., 2016); our current study adds to the literature that glutamatergic signaling in the GPe is altered in PD.

STN-GPe network function and its dysfunction in the context of PD have been widely studied. Experimental and computational studies suggest that the STN-GPe network is important for the generation and amplification of oscillatory activity (Bevan et al., 2002; Gillies et al., 2002; Terman et al., 2002; Walters et al., 2007; Mallet et al., 2008; Holgado et al., 2010; Cruz et al., 2011; Tachibana et al., 2011). Abnormally synchronized beta oscillations (i.e., 15– 30 Hz) in the STN-GPe network are thought to be partially responsible for the hypokinetic symptoms of PD. Abolishing the pathological oscillatory activity by lesioning or deep-brain stimulation of the STN or the GPe has profound therapeutic benefits in alleviating motor symptoms of PD (Albin et al., 1989; Bergman et al., 1990; Hurtado et al., 1999; Vitek et al., 2004; Kuhn et al., 2006; Hammond et al., 2007; Johnson et al., 2012; Vitek et al., 2012; McGregor and Nelson, 2019). We previously observed strengthening of the GABAergic GPe input to the STN with chronic 6-OHDA lesion (Fan et al., 2012); it is now clear that this input arises from PV^+^ neurons as PV^+^ neurons are the primary source of inhibitory input to the STN (Mastro et al., 2014; Hernandez et al., 2015; Abecassis et al., 2020). As the activity of the STN negatively regulates motor output (Hamani et al., 2004; Aron and Poldrack, 2006; Aron et al., 2007; Eagle et al., 2008; Schmidt et al., 2013; Schweizer et al., 2014; Fife et al., 2017; Wessel and Aron, 2017; Adam et al., 2020), a decrease in the ambient glutamate content in the GPe (Cui et al., 2016), along with a reduction in the STN-PV^+^ input, would disinhibit the STN and suppress motor output in the parkinsonian state, thus contributing to the hypokinetic symptoms of PD. On the other hand, a strengthening of the inhibitory PV^+^ input to the STN (Chu et al., 2015) would promote movement and may act as a compensatory mechanism against the hypokinetic effects of the abnormal glutamatergic signaling in the GPe in PD. How these synaptic adaptations interact will vary depending upon on-going activity of the network and the interplay of synaptic features unique to each synapse. As we attempt to make predictions based on rate-based models, we should begin to incorporate oscillation theories with our thinking in order to gain critical insights into basal ganglia function and dysfunction (Wichmann et al., 2011; Wilson, 2013; Little and Brown, 2014).

## Supporting information

Movie 1. In vivo optogenetic stimulation of PV+ neurons promoted locomotion.

Movie 2. In vivo light delivery in PV-Cre mice transduced with a control virus (eYFP only) did not affect locomotion.

Movie 3. In vivo optogenetic inhibition of STN neurons promoted locomotion.

Movie 4. In vivo optogenetic stimulation of Npas1+ neurons suppressed locomotion.

Movie 5. In vivo light delivery in Npas1-Cre-tdTom mice transduced with a control virus (eYFP only) did not affect locomotion.

## Acknowledgments

The authors would like to thank Drs. Daniel Dombeck and Geoffrey Swanson for critical feedback of the work, Dr. Kimberly Ritola for AAVretro production, members of the Chan lab for technical assistance and intellectual exchange, Ayfer Pamukcu and Noyan Pamukcu for their encouragement and compassion, Cooper Chan and Cassidy Chan for their company and emotional support during the COVID-19 shelter-in-place. This work is supported by NIH R01 NS069777 (CSC), P50 NS047085 (CSC), R01 MH112768 (CSC), R01 NS097901 (CSC), R01 MH109466 (CSC), R01 NS088528 (CSC), R00 MH109569 (TNL), DP2 MH122401 (TNL), T32 AG020506 (AP), T32 NS041234 (HSX), and F32 NS098793 (HSX).

## Author contributions

AP conceived the study. AP and QC designed and conducted the electrophysiological measurements. AP and QC designed and conducted the behavioral experiments. EA performed pilot behavioral experiments. HSX performed pilot experiments on the anatomical properties of the STN-GPe input. AP wrote the analysis scripts for *in vivo* and *ex vivo* experiments. AP, BLB, IF, and SC performed histological analysis. AWH, TNL, and SMB offered specialized reagents and technical expertise. AP and CSC wrote the manuscript with input from all co-authors. CSC designed, directed, and supervised the project. All authors reviewed and edited the manuscript.

## Movie Legends

Movie 1. *In vivo* optogenetic stimulation of PV^+^ neurons promoted locomotion.

Representative movie of a mouse in an open field arena showing increased locomotion upon optogenetic stimulation of PV^+^ neurons. Behavior from two consecutive trials are shown. Video is shown at 2x speed.

Movie 2. *In vivo* light delivery in PV-Cre mice transduced with a control virus (eYFP only) did not affect locomotion.

Representative movie of a mouse in an open field arena showing the lack of effect in locomotion upon light delivery to PV^+^ neurons with no opsin expression. Behavior from two consecutive trials are shown. Video is shown at 2x speed.

Movie 3. *In vivo* optogenetic inhibition of STN neurons promoted locomotion.

Representative movie of a mouse in an open field arena showing increased locomotion upon optogenetic inhibition of STN neurons. Behavior from two consecutive trials are shown. Video is shown at 2x speed.

Movie 4. *In vivo* optogenetic stimulation of Npas1^+^ neurons suppressed locomotion.

Representative movie of a mouse in an open field arena showing decreased locomotion upon optogenetic stimulation of Npas1^+^ neurons. Behavior from two consecutive trials are shown. Video is shown at 2x speed.

Movie 5. *In vivo* light delivery in Npas1-Cre-tdTom mice transduced with a control virus (eYFP only) did not affect locomotion.

Representative movie of a mouse in an open field arena showing the lack of effect in locomotion upon light delivery to Npas1^+^ neurons with no opsin expression. Behavior from two consecutive trials are shown. Video is shown at 2x speed.

